# Multiple ribonuclease A family members cleave transfer RNAs in response to stress

**DOI:** 10.1101/811174

**Authors:** Yasutoshi Akiyama, Shawn Lyons, Marta M. Fay, Takaaki Abe, Paul Anderson, Pavel Ivanov

## Abstract

During stress, changes in gene expression are critical for cell survival. Stress-induced tRNA cleavage has been implicated in various cellular processes, where tRNA fragments play diverse regulatory roles. Angiogenin (ANG), a member of the RNase A superfamily, induces cleavage of tRNAs resulting in the formation of tRNA-derived stress-induced RNAs (tiRNAs) that contribute to translational reprogramming aiming at cell survival. The role of other stress-induced RNases in tRNA cleavage is poorly understood. Using gene editing and biochemical approaches, we show that other members of the RNase A family are capable of targeting tRNAs in stress-responsive manner. We show that in the absence of ANG, these RNases also promote the production of tiRNAs. Moreover, specific stresses (such as treatment with sodium arsenite) activate cleavage of universal 3’-CCA termini of tRNAs in ANG-independent fashion in living cells. We conclude that multiple RNase A family members are capable of targeting tRNAs in a stress-specific manner *in vivo*.

## INTRODUCTION

Angiogenin (ANG) is a secreted ribonuclease (RNase) that is a member of the RNase A superfamily (35% amino acid identity to RNase A) (1). Although, ribonucleolytic activity of ANG is only a fraction of RNase A (2), most of its biological functions are critically dependent on its RNase activity (3-5). Although first identified as a tumour angiogenesis factor, ANG has been implicated in tumorigenesis, neurodegeneration, inflammation, pregnancy and innate immunity (reviewed in (6)).

ANG is a stress-responsive RNase that is transcriptionally upregulated by stress (7,8) and modulates its RNase activity in response to stresses, especially oxidative stress (reviewed in (9)). Under optimal growth conditions, ANG is predominantly located in the nucleus where it stimulates rRNA biogenesis to promote cell growth and proliferation. While a small proportion of ANG is found in the cytoplasm, it is held in an inactive state by its inhibitor, RNH1. In response to different stress stimuli, ANG translocates into the cytoplasm from the nucleus and dissociates from RNH1 (10). Under these conditions, ANG cleaves cytoplasmic tRNAs within their anticodon loops, generating two smaller RNA species called tRNA-derived stress-induced RNAs (tiRNAs) (11) or tRNA halves (12). The functional role of 5’- and 3’-tiRNAs are just beginning to be determined. 5’-tiRNAs derived from tRNA^Ala^ and tRNA^Cys^ inhibit translation initiation by displacing the eIF4F complex from the cap structures of mRNAs (13) and facilitate the formation of stress granules (SGs), pro-survival cytoplasmic foci containing stalled pre-initiation ribosomal complexes (14-17). In addition to translational repression, tiRNAs also promote survival by binding to Cytochrome *c* to prevent apoptosome formation (18). Through these mechanisms, ANG promotes cell survival at low metabolic cost under stress conditions.

*In vitro*, ANG efficiently cleaves 5S ribosomal RNA (19) and other small RNAs such as snRNAs and tRNAs (20). *In vivo*, its enzymatic activity appears to be specific and limited to tRNAs involved in translation (21), and promoter-associated RNAs, which modulate ribosomal DNA (rDNA) transcription (22). *In vitro*, ANG targets not only the anticodon but also the 3’-CCA terminus of tRNAs (23), the triplet that is added post-transcriptionally during tRNA maturation by the CCA-adding enzyme TRNT1. CCA addition is required for the maturation of both nuclear and mitochondrial tRNAs, and tRNAs without CCA ends are translationally-incompetent. Partial loss-of-function mutations in the TRNT1 are linked to the congenital sideroblastic anemias, disorders originating from metabolic effects in both cytosol and mitochondria (24). To date, anticodons of tRNAs and promoter-associated RNAs are known as genuine substrates *in vivo*.

The same study showed that oxidative stress induced by sodium arsenite (SA), known inducer of ANG-mediated tRNA cleavage, also promotes CCA removal from tRNAs in live cells (23). Removal of the terminal adenine residue from the 3’-CCA by ANG can be reversed *in vitro* by recombinant TRNT1, thus making tRNAs chargeable again. Authors have hypothesized that such reversible mechanism would also act as a stress-responsive switch to repress and reactivate translation at a low metabolic cost in live cells. However, since this model is based largely on *in vitro* experiments, whether it is valid *in vivo* is still unclear.

Here, we directly examined whether ANG targets 3’-CCA termini of tRNAs *in vivo* using high-throughput RNA sequencing and biochemical approaches based on RNA ligation that we developed. We determined that although ANG targets CCA ends *in vitro*, it primarily targets anticodon loops and not CCA ends in live cells. However, and in agreement with data from the Ignatova lab, SA promotes cleavage of the CCA ends, although in an ANG-independent manner. This SA-induced cleavage is regulated by RNH1, a universal inhibitor of ribonucleases belonging to the RNase A superfamily, suggesting that other RNases related to ANG are also stress-responsive and tRNA specific. Moreover, our data also suggest that in the absence of ANG, other RNases are capable to generate tiRNAs providing “compensatory” tRNA cleavage in stressed cells.

## MATERIAL AND METHODS

### Cell culture and treatment

U2OS cells were cultured at 37°C in a CO_2_ incubator in Dulbecco’s modified Eagle’s medium (DMEM) supplemented with 10% fetal bovine serum (FBS) (Sigma) and 1% of penicillin/streptomycin (Sigma). For ANG treatment, U2OS cells were incubated with DMEM supplemented with 0.5 µg/ml recombinant human ANG for 1 hour. Human recombinant ANG was prepared as reported previously (25).

### CCA-specific ligation

Total RNAs were incubated at 37°C for 40 min in 20 mM Tris-HCl (pH 9.0) for deacylation of mature tRNAs, followed by purification with RNA Clean & Concentrator Kits (Zymo Research). Ten micrograms of deacylated total RNAs were incubated with 400 pmol of hairpin oligo(H-oligo) or double-strand oligo (ds-oligo) at 90°C for 2 min for denaturing. Denatured RNAs were then incubated in 5 mM Tris-HCl (pH 8.0), 0.5 mM EDTA and 10 mM MgCl_2_ at 37°C for 15 min in 50-µl reaction volume for annealing. For RNA ligase 2-based ligation reaction, the annealed samples were incubated with 5 µl of 1x reaction buffer, 5 U (0.5 µl) of T4 RNA ligase 2 (Rnl2) (New England Biolabs) and 40 U (1 µl) of RNasin (Promega) at 37°C for 1 hr, followed by overnight incubation at 4°C. For DNA ligase-based ligation, annealing was carried out in 16-µl reaction volume. Annealed samples were then incubated with 2 µl of 1x reaction buffer, 400 U (2 µl) of T4 DNA ligase (New England Biolabs) and 40 U (1 µl) of RNasin overnight at 16°C. The sequences of oligos were shown in Supplementary Table 1.

### CC-specific ligation combined with PNK pre-treatment

The sequences of oligos for CC-specific ligation were shown in Supplementary Table 1. Before ligation reaction, samples were pre-treated with 10 U of T4 polynucleotide kinase (PNK) (New England Biolabs) at 37°C for 1 hr according to the manufacturer’s instruction to remove 2’, 3’-cyclic phosphate. Ligation reaction was performed with Rnl2 in the same way as Ligation 3 in Figure 3A.

**Figure 1.**
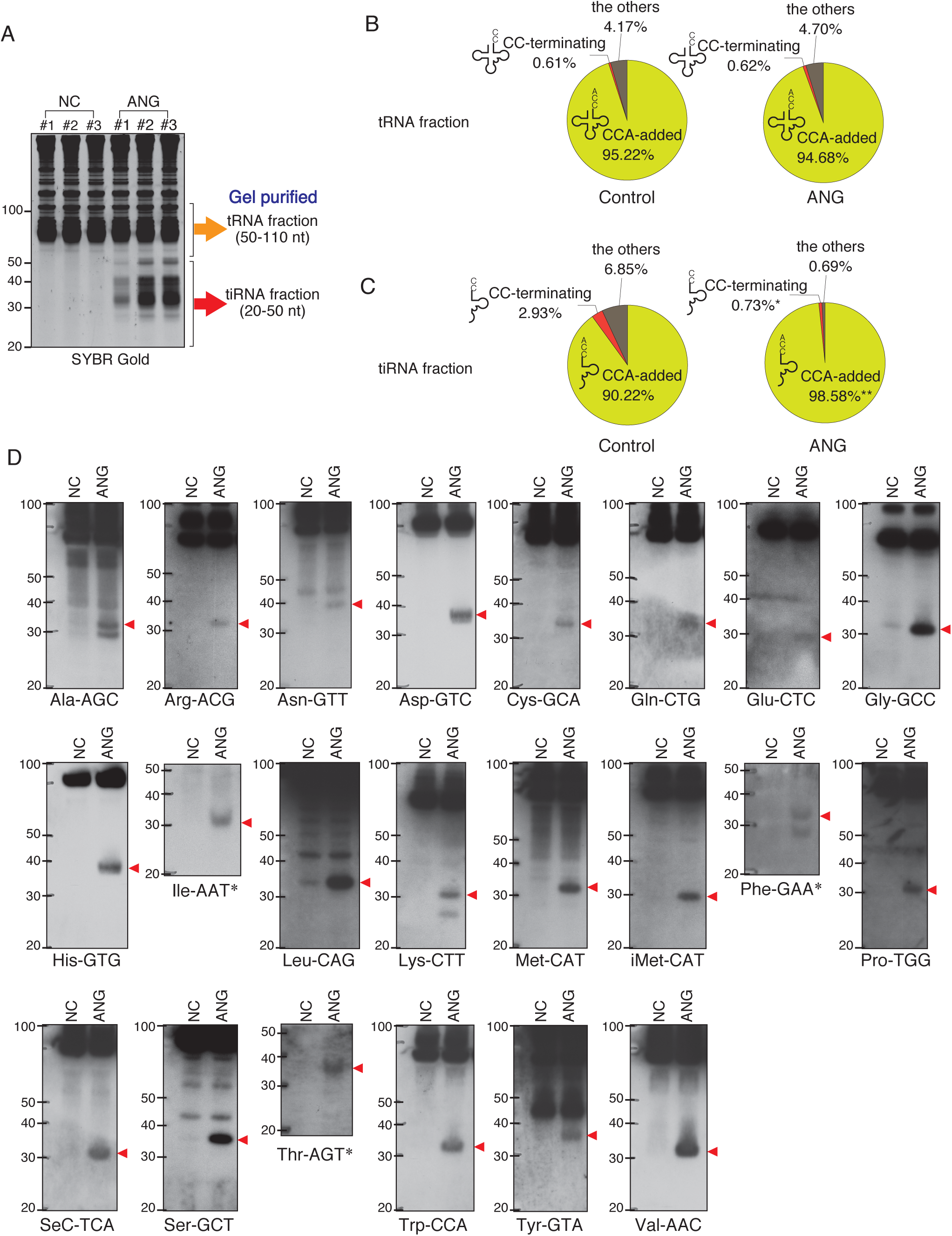
Effect of ANG treatment on the proportions of the tRNA-derived reads that have 3’-CCA or 3’-CC termini. (A) ANG-mediated tiRNA production in U2OS cells detected by SYBR Gold staining. The libraries were generated from gel-purified tRNA fraction (50-110 nt) or tiRNA fraction (20-50 nt). (B) Distribution of reads (in %) in tRNA fraction. Raw data can be found in table S2A. (B) Distribution of reads (in %) in tiRNA fraction. Raw data can be found in Supplementary Table 1. The results are shown as pie charts. *: p<0.05 VS Control, **: p<0.01 VS Control

**Figure 2.**
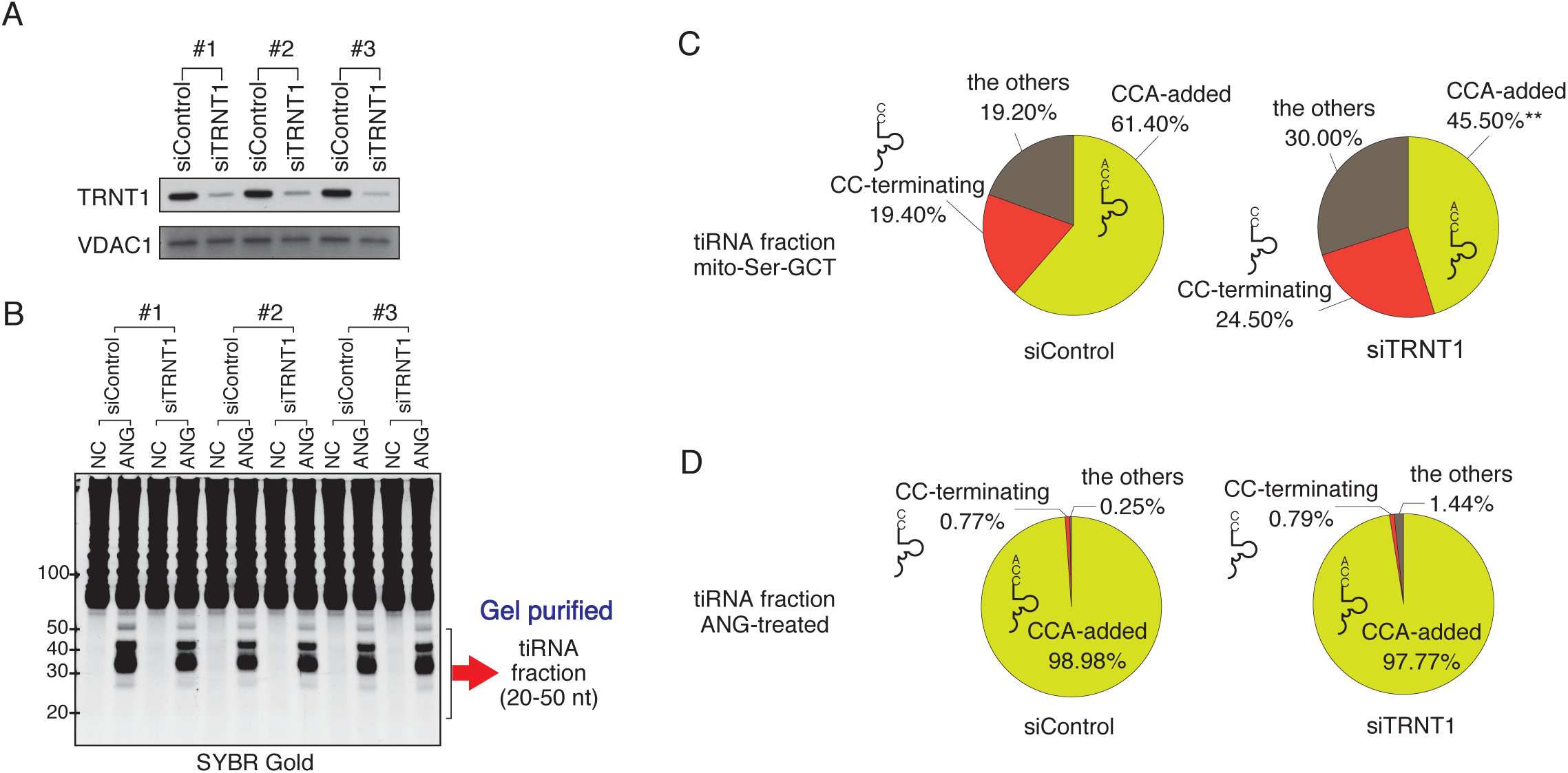
Effect of TRNT1 depletion on the proportion of tRNA-derived reads with 3’-CCA termini. (A) Knockdown efficiency of TRNT1 (siTRNT1, 96 h post-transfection) was evaluated by Western blot using TRNT1-specific antibodies. Control siRNA (siControl) was used as control. VDAC1 was used as a loading control. Three biological replicates are shown. (B) SYBR Gold staining of total RNA prepared from cells treated with control or TRNT1-specific siRNAs (as shown in Figure 4A) and left untreated (NC) or treated with recombinant ANG (ANG). tiRNA fraction (20-50 nt) was gel-purified and used for library preparation. (C-D) Effect of TRNT1 knockdown on the proportions of 3’-tiRNAs that have 3’-CCA or 3’-CC termini. (C) Distribution of reads (in %) in mitochondrial tRNA-Ser-GCT and (D) nuclear-encoded tRNAs. Raw data can be found in Supplementary Table 2.

**Figure 3.**
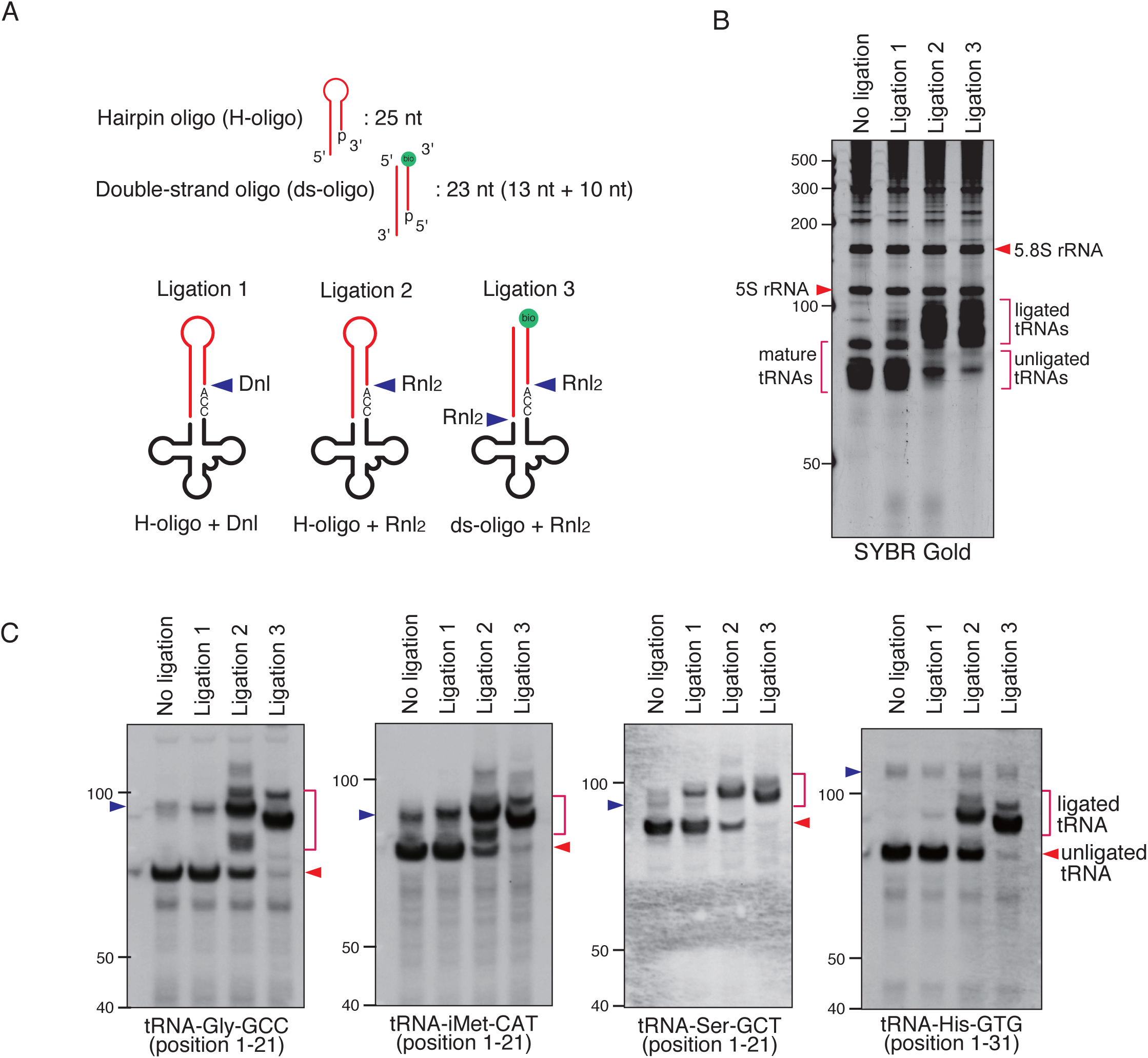
Validation of CCA-specific ligation methods. (A) Schema for three CCA-specific ligation methods. Dnl: T4 DNA ligase, Rnl2: T4 RNA ligase 2, bio: biotin. (B-C) The method using double-strand oligo and RNA ligase 2 has the best ligation efficiency. (B) SYBR Gold staining and (C) Northern blotting for CCA-specific ligation products. The blue arrowheads indicate the bands for pre-tRNAs.

### Northern blotting

Total RNA was extracted by using Trizol (Invitrogen). RNA was run on 10% or 15% TBE-urea gels (ThermoFisher Scientific), transferred to positively charged nylon membranes (Roche). The membranes were cross-linked by UV irradiation except for detecting 5’-tiRNA-Ile-AAT, -Phe-GAA and -Thr-AGT. For detecting them, chemical cross-linking using 1-ethyl-3-(3-dimethylaminopropyl) carbodiimide (EDC, Sigma) was performed to improve the sensitivity as previously reported (26). After cross-linking, the membranes were hybridized overnight at 40°C with digoxigenin (DIG)-labeled DNA probes in DIG Easy Hyb solution (Roche). After low stringency washes (washing twice with 2× SSC/0.1% SDS at room temperature) and high stringency wash (washing once with 1× SSC/0.1% SDS at 40°C), the membranes were blocked in blocking reagent (Roche) for 30 min at room temperature, probed with alkaline phosphatase-labeled anti-digoxigenin antibody (Roche) for 30 min, and washed with 1x TBS-T. Signals were visualized with CDP-Star ready-to-use (Roche) and detected using ChemiDoc imaging system (BioRad) according to the manufacturer’s instructions. Oligonucleotide probes were synthesized by IDT. DIG-labeled probes were prepared using the DIG Oligonucleotide tailing kit (2nd generation; Roche) according to the manufacturer’s instructions. The sequences of the probes were shown in Supplementary Table 2.

### Generation of ANG and RNH1 knockout cells

ANG knockout and RNH1 knockout U2OS cells were generated using CRISPR/Cas9-mediated gene editing as previously reported (27). Briefly, oligonucleotides corresponding to a gRNA targeting the sequence 5’-TGGTTTGGCATCATAGTGCT-3’ in ANG and 5’-GAGCCTGGACATCCAGTGTG-3’ in RNH1 were designed using CRISPR Design software from the Zhang lab (crispr.mit.edu). Oligonucleotides were annealed and cloned into the pCas-Guide vector (Origene) according to the manufacturer’s protocol, and the resulting plasmid was co-transfected with pDonor-D09 (GeneCopoeia), which carries a puromycin resistance cassette, into U2OS cells using Lipofectamine 2000 (Invitrogen). The following day, cells were selected with 1.5 μg/ml of puromycin for 24 hours only, to lessen the likelihood of genomic incorporation of pDonor-D09. Cells were cloned by limiting dilution and screened by western blot analysis using anti-ANG antibody (Santa Cruz, C-1) and anti-RNH1 antibody (Proteintech group, 10345-1-AP). Anti-beta-actin antibody (Proteintech group, 66009-1-Ig) was used as a loading control. Knockout was confirmed by genotyping (Fig. S5).

### Knockdown of TRNT1

U2OS cells were transfected with siRNA against TRNT1 (ON-TARGETplus SMARTpool; GE Dharmacon, catalog #L-015850-00-0005) or Control siRNA (ON-TARGETplus Non-targeting siRNA #1; GE Dharmacon, catalog #D-001810-01-05) at a concentration of 40 nM using Lipofectamine 2000 (Invitrogen) according to the manufacturer’s protocol for reverse transfection. Ninety-six hours after transfection, cells were subjected to ANG treatment described above. Knockdown efficiency was checked by western blot analysis using anti-TRNT1 antibody (Novus, NBP1-86589). Anti-VDAC1 antibody (Santa Cruz, N-18) was used as a loading control.

### *In vitro* ANG digestion

*In vitro* ANG digestion was performed according to the procedure described in (23) with slight modification. Briefly, total RNA in 30 mM HEPES (pH 7.0), 30 mM NaCl was heated at 90°C for 2 min and cooled down to room temperature. MgCl_2_ and BSA were added to final concentrations of 2 mM and 0.01%, respectively, and further incubated at 37°C for 5 min. Recombinant human ANG was added to a final concentration of 0.2 µM and incubated at 37°C for 4 hours.

### RNA sequencing library preparation for high-throughput sequencing

RNA-seq libraries were prepared from samples from three independent experiments. 10 µg of total RNA was run on 15% Urea-TBE gel, and 20-50 nt (tiRNA fraction) or 50-110 nt (tRNA fraction) was gel-purified using ZR small-RNA PAGE Recovery kit (Zymo Research). The purified RNAs were treated with calf intestinal phosphatase (New England Biolabs). After purification using Direct-zol RNA Microprep (Zymo Research), the RNAs were treated with T4 polynucleotide kinase (New England Biolabs), and then purified using Direct-zol RNA Microprep. Small RNA libraries were prepared using the TruSeq Small RNA library preparation kit (Illumina) according to the manufacturer’s protocol. Sequencing was performed on the Illumina platform (Molecular Biology Core Facility, Dana-Farber Cancer Institute, Boston, USA), and 75 bp single-end reads (tiRNA fraction) or 150 bp single-end reads (tRNA fraction) were generated.

### Calculation of the proportion of fragments that have CCA or CC at their 3’-termini

For accurate mapping to the reference genome, we generated two sets of adapter-trimmed data using cutadapt (28). In the first set, only the adapter sequence was trimmed (“Only Adapter-trimmed”), and in the second, the 3’-CCA or 3’-CC and adapter sequences were trimmed (“CCA-trimmed”) (Supplementary Figure 3B). Then “CCA-trimmed” data set was aligned to hg38 reference genome (obtained from UCSC Genome Browser (29)) using Bowtie (30) with parameters “-k 1 --best -v 3” which reports one best match with up to 3 mismatches. Up to 3 mismatches were allowed in order to prevent mapping failure for tRNA-derived reads with misincorporations due to modification sites. Reads that overlapped 3’-end (5 nt) of tRNA genes were extracted as “3’-end containing reads” using samtools (31) (Supplementary Figure 3C). The reads with the same ID were extracted as FASTQ format files from “Only Adapter-trimmed” data using seqtk (https://github.com/lh3/seqtk). There are 109 tRNA genes and 22 tRNA genes ending with CCA and CC, respectively (Supplementary Table 3D). Immature tRNAs (end-processed, but without CCA addition) derived from these genes also ends with CCA or CC. To distinguish these reads from CCA-added or CC-terminating reads, 631 tRNA genes were divided into 3 groups: 1) ending with CCA (109 genes), 2) ending with CC (22 genes), and 3) the others (500 genes). Then the number and the proportions of CCA-added or CC-terminating reads were calculated in each group (Supplementary Table 3D). The annotation information of genomic (cytoplasmic) tRNA genes were obtained from UCSC Genome Browser (29). Genomic tRNA sequences were obtained from genomic tRNA database (gtRNAdb) (32). The sequence and annotation information of mitochondrial tRNA-Ser-GCT were obtained from the Mamit-tRNA database (33). The read counts were presented as reads per million mapped reads (PMMR).

### Statistical analyses

Comparisons between groups were performed using one-way ANOVA or an unpaired Student’s t-test as appropriate. The Tukey–Kramer test was used for multiple comparison.

## RESULTS

### Angiogenin does not target CCA ends of tRNAs *in vivo*

ANG is reported to target tRNAs in anticodon loops and CCA ends *in vitro* (23). To clarify directly the effect of ANG on tRNA 3’-CCA termini *in vivo*, we determined whether ANG treatment changed the proportion of CCA-added tRNAs or 3’-tiRNAs using high-throughput RNA sequencing (RNA-seq). Since it has been reported that ANG cleaves within the 3’-CCA termini between the penultimate C and terminal A (23), if a similar reaction occurs *in vivo*, cleaved tRNAs should bear CC-termini. Therefore, we calculated the number and proportion of tRNAs or 3’-tiRNAs with CC as well as with CCA at their 3’-termini.

Recombinant ANG added to cell culture media is rapidly internalized and cleaves mature cytoplasmic tRNAs (11,34). We verified that ANG treatment induced tiRNA production (Fig. 1A, 3 biological replicates). Next, we generated RNA sequencing libraries from gel-purified fractions (tRNA fraction and tiRNA fraction, Fig. 1A). ANG-cleaved 5’-tRNA products start with 5’-monophosphates and end with 2’, 3’-cyclic phosphates, whereas 3’-tRNA products start with a hydroxyl residue and end with a hydroxyl residue (35). Since traditional RNA-seq libraries are designed to efficiently capture 5’-monophosphate/3’-hydroxyl groups of RNAs, ANG-derived RNA fragments are not optimal for library construction. Therefore, we treated total RNA samples with calf intestinal phosphatase (CIP) and T4 polynucleotide kinase (PNK) prior to library preparation in order to convert tiRNAs to proper substrates for library preparation (Fig. S1A). Using Illumina sequencing, we then generated 75 bp single-end reads (tiRNA fraction) or 150 bp single-end reads (tRNA fraction) and calculated the proportion of these reads that end with CCA or CC (as described in Fig. S1A-D, and table S3).

In the tRNA fraction, the total number of reads (mainly mature tRNAs) was slightly but significantly decreased by ANG treatment (188,990 PMMR (per million mapped reads) and 164,810 PMMR for control and ANG, respectively) (Table S3A). CCA-terminating tRNAs were also slightly decreased by ANG treatment (179,947 PMMR and 156,044 PMMR for control and ANG, respectively). However, the number of CC-terminating tRNAs was not increased by ANG treatment (1,170 PMMR and 1,020 PMMR for control and ANG, respectively, Table S3A), suggesting that the decrease of CCA-added tRNAs was not due to the ANG-mediated cleavage of 3’-CCA termini, but due to the decrease of full length tRNAs by ANG-mediated tiRNA production by targeting anticodon loops. The proportion of tRNAs with CCA did not change significantly following ANG treatment, 95.22% for control and 94.68% for ANG (Fig. 1B and Table S3A). The proportion of CC-terminating tRNAs was small and not altered by ANG treatment, 0.61% for control and 0.62% for ANG (Fig. 1B and table S3A). These data demonstrate that ANG does not efficiently target tRNA 3’-CCA termini *in vivo*, as most the tRNAs remain intact and unchanged after ANG treatment.

It has been reported that 3’-CCA cleavage by ANG precedes tiRNA production *in vitro* (23). If this occurs *in vivo*, most 3’-tiRNAs would be CCA-cleaved. We examined the 3’-termini of 3’-tiRNAs after treating cells with ANG (Table S3B and Fig. 1C). ANG treatment results in the production of 5’-tiRNAs from all tRNA species (represented by 22 amino acids), thus reassuring that potential 3’-CCA cleavage step has been already occurred (23) (Fig. 1D). The total number of reads (mostly 3’-tiRNAs) was significantly increased by ANG treatment (from 38,375 PMMR in control to 240,173 PMMR in ANG, ∼6.3-fold increase) consistent with previous reports (11,12,36). Further analysis indicated that most 3’-tiRNAs (98.58%) have CCA at their 3’-termini (Fig 1C and Table S3B). Furthermore, the number of CC-terminating 3’-tiRNAs was not significantly increased by ANG treatment (1,151 PMMR and 1,765 PMMR for control and ANG, respectively). On the other hand, %CC was significantly decreased by ANG treatment (2.93% in control to 0.73% in ANG) due to the marked increase of CCA-added (i.e. intact) 3’-tiRNAs (Fig. 1C).

### TRNT1 does not repair CCA-ends *in vivo* after ANG cleavage

*In vitro*, TRNT1 can repair CCA ends after ANG-mediated cleavage (23). In order to assess whether a similar mechanism occurs *in vivo*, we examined whether the CCA-adding enzyme TRNT1 is involved in CCA repair following ANG treatment. To do so, we used siRNAs targeting TRNT1. We reproducibly achieved an approximately 85% knockdown of TRNT1 protein (Fig. 2A). RNAi mediated knockdown of TRNT1 caused no detectable difference in ANG-mediated tiRNA production (Fig. 2B). Under these conditions, we generated RNA-seq libraries using gel-purified tiRNA fractions. To further confirm that TRNT1 knockdown affected TRNT1 activity, we calculated %CCA of mitochondrial tRNA-Ser-GCT, because mitochondrial tRNA-Ser-GCT is most susceptible to TRNT1 hypofunction (37). In patients with TRNT1 hypomorphic mutations, CCA addition to mitochondrial tRNA-Ser-GCT is significantly affected (37). As expected, the total number of 3’-fragment reads derived from mitochondrial tRNA-Ser-GCT was significantly decreased by ANG treatment (59.0 PMMR and 21.4 PMMR for siControl-NC and siControl-ANG, respectively. p=0.0068), because of the marked increase of tiRNAs derived from cytoplasmic tRNAs. However, ANG treatment did not induce significant cleavage of mitochondrial tRNA-Ser-GCT (Fig. 2C, Table S4). In the non-treated samples (NC column, Table S4), the number of CCA-added 3’-fragments was significantly decreased by TRNT1 knockdown (36.6 PMMR and 17.5 PMMR for NC-siControl and NC-siTRNT1, respectively, p=0.0409). The proportion of CCA-added 3’-fragments was also significantly decreased by TRNT1 knockdown (61.4% and 45.5% for NC-siControl and NC-siTRNT1, respectively. p=0.0066). These data demonstrate that TRNT1 knockdown was effective enough to have a functional impact on CCA addition.

We then calculated the proportion of CCA-added and CC-terminating 3’-tiRNAs derived from cytoplasmic tRNAs in control and TRNT1 knockdown cells treated or not treated with ANG (Fig. 2D and Table S4). As expected, ANG treatment increased levels of 3’-tiRNAs significantly both in control and TRNT1 knockdown cells. However, the percentage of CCA-terminating cytoplasmic 3’-tiRNAs did not change significantly following TRNT1 knockdown (98.98% and 97.77% for siControl and siTRNT1, respectively; p=0.9731). In addition, their percentage was not affected by TRNT1 knockdown in ANG-treated cells (0.77% and 0.79% for siControl and siTRNT1, respectively. p=0.9998) (Figure 2D and Table S4). These results show that TRNT1 does not play a role in regulating ANG-dependent cleavage of tRNAs *in vivo*.

### Probing integrity of tRNAs’ CCA-ends *in vivo* using cyclic phosphate specific ligation method

Because tRNA has a stable structure and is heavily decorated with post-transcriptional modifications, our tRNA sequencing results can be biased. As alternative approach, we directly purified tiRNAs from U2OS cells treated with ANG, and biochemically analysed them. As ANG leaves 2’, 3’-cyclic phosphate at the 3’ end of the 5’-fragment after anti-codon cleavage (16,38,39), we took advantage of the RNA ligase RtcB which specifically ligates 2’, 3’-cyclic phosphate residues to 5’-hydroxyls (40). We reasoned that RtcB will specifically recognize ANG-cleaved 5’-fragments containing cyclic phosphates at their 3’ends and ligate them to a synthetic oligo with a 5’-hydroxyl (Fig. S2). First, we treated total RNA isolated from osteosarcoma U2OS cells with recombinant ANG *in vitro*, and gel-purified representative tiRNA fractions (Fig. S3A). Using recombinant *E.coli* RtcB ligase, we then performed ligation reactions between synthetic 5’-hydroxyl, 3’-biotinylated oligo and tiRNA fractions (Fig. S3A-C). As a positive control, we performed a reaction using purified endogenous 5’-tiRNA (5’-tiRNA) and endogenous 3’-tiRNA (3’-tiRNA) fractions. RtcB ligated 5’-tiRNAs to 3’-tiRNAs to produce full length (i.e. repaired) tRNAs (Fig. S3B). 3’-tiRNAs and 5’-tiRNAs were also ligated to the synthetic oligo and the ligation products were detected by both SYBR Gold staining (Fig. S3B) and streptavidin-HRP blotting (Fig. S3C). The length of ligated products was compatible with the predicted ligation product (“5’-tiRNA + 3’-tiRNA” > “3’-tiRNA + oligo” > “5’-tiRNA + oligo”) (Fig. S3K).

Next, we performed similar reactions using the gel-purified tRNA fraction. We reasoned that if ANG cleaves CCA ends, then ANG-treated tRNAs will be appropriate substrates for RtcB-mediated ligation. Indeed, the reaction between ANG-treated tRNAs and the synthetic oligo with RtcB generated a band around 100 nt, that was detected by both SYBR Gold staining (Fig S3D) and streptavidin-HRP blotting (Fig. S3E). The length of the ligation product was compatible with the predicted length of tRNA + oligo (Fig. S3K). Thus, in agreement with published data (23), ANG targets tRNA CCA ends *in vitro*.

To clarify the potential difference between ANG-mediated cleavage *in vivo* and *in vitro*, we performed the same experiments using the tiRNA fractions (Fig. S3F) and tRNA fractions (Fig. S3I) produced by *in vivo* treatment of U2OS cells with recombinant ANG. Reactions between purified endogenous 5’-tiRNA (5’-tiRNA) and endogenous 3’-tiRNA (3’-tiRNA) fractions served as control (Fig. S3G). Although not detected by SYBR Gold staining (Fig. S3G), 5’-tiRNAs were ligated to the synthetic oligo and the ligation product was detected by blotting and probing with streptavidin-HRP (Fig. S3H). This ligation product was shorter than that derived from the reaction between 5’-tiRNA and 3’-tiRNA detected in Fig 1G, which is compatible with the predicted length of the ligation products (Fig. S3K). On the other hand, no ligation product was detected by the reaction between the synthetic oligo and 3’-tiRNA (Fig. S3G and S3H). Together these data indicate that only endogenous 5’ tiRNAs, which contain a 2’, 3’-cyclic phosphate, are substrates for RtcB-mediated ligation. In addition, no ligation product was detected in the reaction of the synthetic oligo and tRNAs derived from ANG-treated cells by streptavidin-HRP (Fig. S3I-J). These data suggest that CCA ends of neither 3’-tiRNAs nor mature tRNAs are cleaved by ANG *in vivo*.

### Probing integrity of tRNAs’ CCA-ends *in vivo* using CCA-specific ligation method

Our ligation using RtcB enzyme is dependent on its ability to ligate RNAs with the 2’-3’-cyclic phosphate moiety. It can be that the SA-induced cleavage of the CCA ends is ANG-/RNase A superfamily member-independent, and thus our RtcB-based approach is inconclusive. To probe this situation, we chose alternative approaches based on the ability of DNA ligase (Dnl) or RNA Ligase 2 (Rnl2) to ligate RNA/DNA chimeric substrates that contain 5’-monophospate. We chose to compare these enzymes to determine their ligation efficiencies since inefficient ligation can strongly impact possible results (41). We also chose two substrates that were previously used to study the integrity of CCA ends, namely hairpin oligo (23,42) and double-stranded oligo (ds-oligo, (43)) (Fig. 3A). As is seen in Fig. 3B-C, the combination of the Rnl2 ligase with ds-oligo results in the most efficient capture and ligation of mature CCA-containing tRNAs as judged by total tRNA shift by SYBR Gold staining (Fig. 3B) and by the northern blotting against individual tRNA species (Fig. 3C).

We used this approach to determine whether SA and/or ANG promote cleavage of the CCA ends *in vivo*. We treated U2OS cells with SA or recombinant ANG, purified total RNA from the cells and proceeded with CCA-specific ligation using Rnl2 and ds-oligo containing biotin (Fig. 3A. ligation 3). Untreated U2OS cells served as a control (Fig. 4A, NC). RNA samples from all conditions were efficiently ligated as judged by SYBR Gold staining (Fig. 4A, CCA-ligation) and immuno-blotting against biotin-containing oligo (Fig. 4B). Using probes against specific tRNA species (tRNA-Gly-GCC and tRNA-iMet-CAT, Fig. 4C), we saw no difference between control and ANG-treated samples but slight yet reproducible decrease of ligation in the SA-induced samples (Fig. 4C, unligated tRNA) suggesting that SA/oxidative stress affects CCA cleavage. It should also be noted that treatment of the purified total RNA with SA at various concentration *in vitro* does not trigger degradation, cleavage or shortening of tRNAs (Fig. S4) thus rejecting the possibility that SA chemically triggers tRNA cleavage. Taken together, these data demonstrate that ANG does not efficiently cleave 3’-CCA termini of tRNAs *in vivo.*

**Figure 4.**
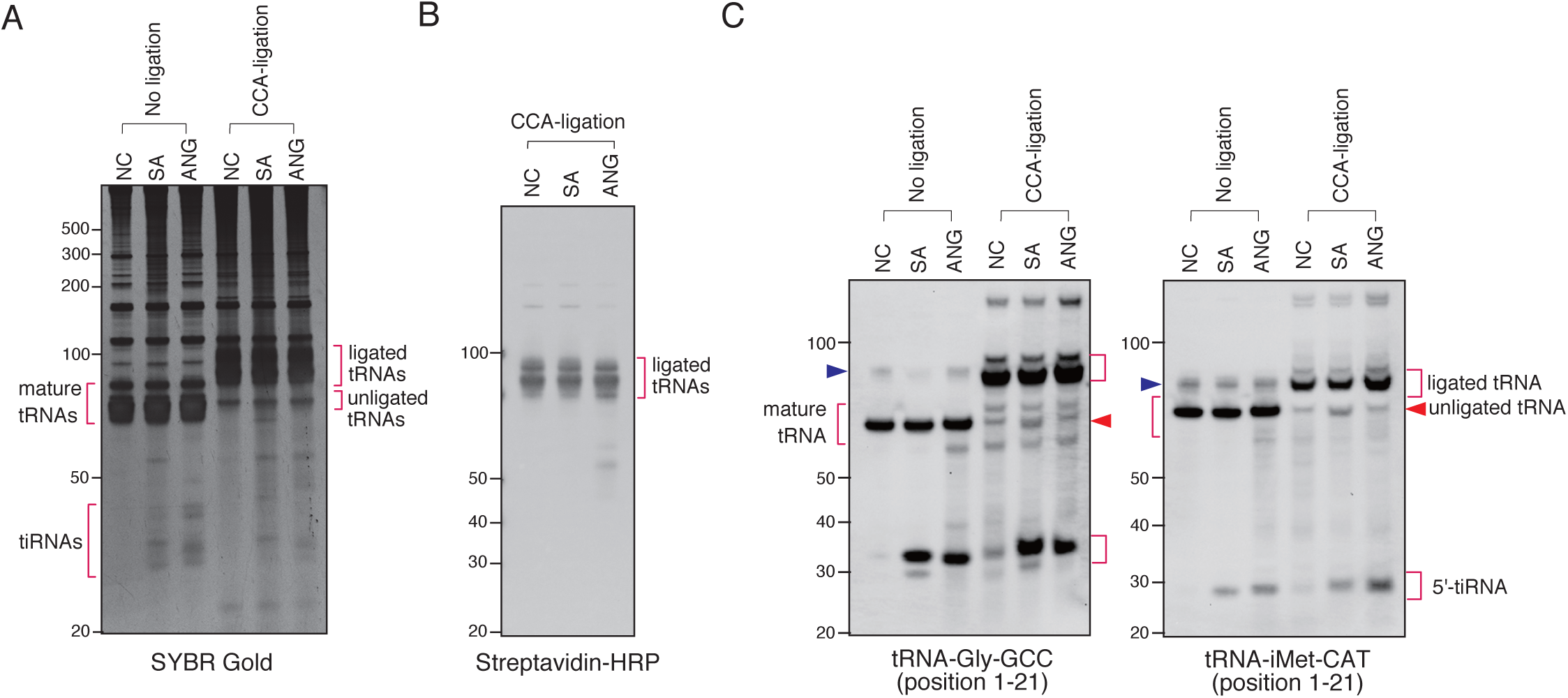
Sodium arsenite treatment induces CCA-deactivated tRNAs. (A) SYBR Gold staining of the CCA-specific ligation products. (B) Visualization of ligated tRNAs by streptavidin-biotin system. (C) Sodium arsenite treatment slightly increases CCA-deactivated tRNAs. The blue arrowheads indicate the bands for pre-tRNAs.

### Sodium arsenite promotes cleavage of tRNAs’ CCA-ends *in vivo*

To complement these findings, we have developed an alternative approach that aims on the ligation of only CC-terminating tRNAs. In this method, ds-oligo has 5’-monophosphate and can only be ligated by Rnl2 to the CC-terminating tRNAs with 3’-hydroxyl group. Thus, if SA induces an RNase that cleaves tRNA CCA ends to produce tRNAs terminating with CC-end with cyclic 2’-3’-phosphate, such substrate will not be ligated (Fig. 5A). As a control, we used treatment with T4 PNK, which removes both a 3’-phosphate and 2’-3’ cyclic phosphate to form a 3’-hydroxyl end. As is seen in Fig. 5B-C, Rnl2 more efficiently ligates tRNAs from SA-treated cells that were pre-treated with T4 PNK suggesting that CC-terminating tRNAs contain 2’-3’ cyclic phosphates at their 3’ends. In turn, this indicates that an enzyme that triggers CCA cleavage under SA stress may belong to the RNase A superfamily members. *In vitro*, SA does not affect the integrity of tRNA CCA ends (Fig. S4). In contrast, *in vitro* digestion of tRNAs leads to the accumulation of tRNAs with CCA-cleaved ends as judged by our CCA-specific ligation (Fig. 6A-B) and the CC-specific ligation (Fig. 6C-D) approaches.

**Figure 5.**
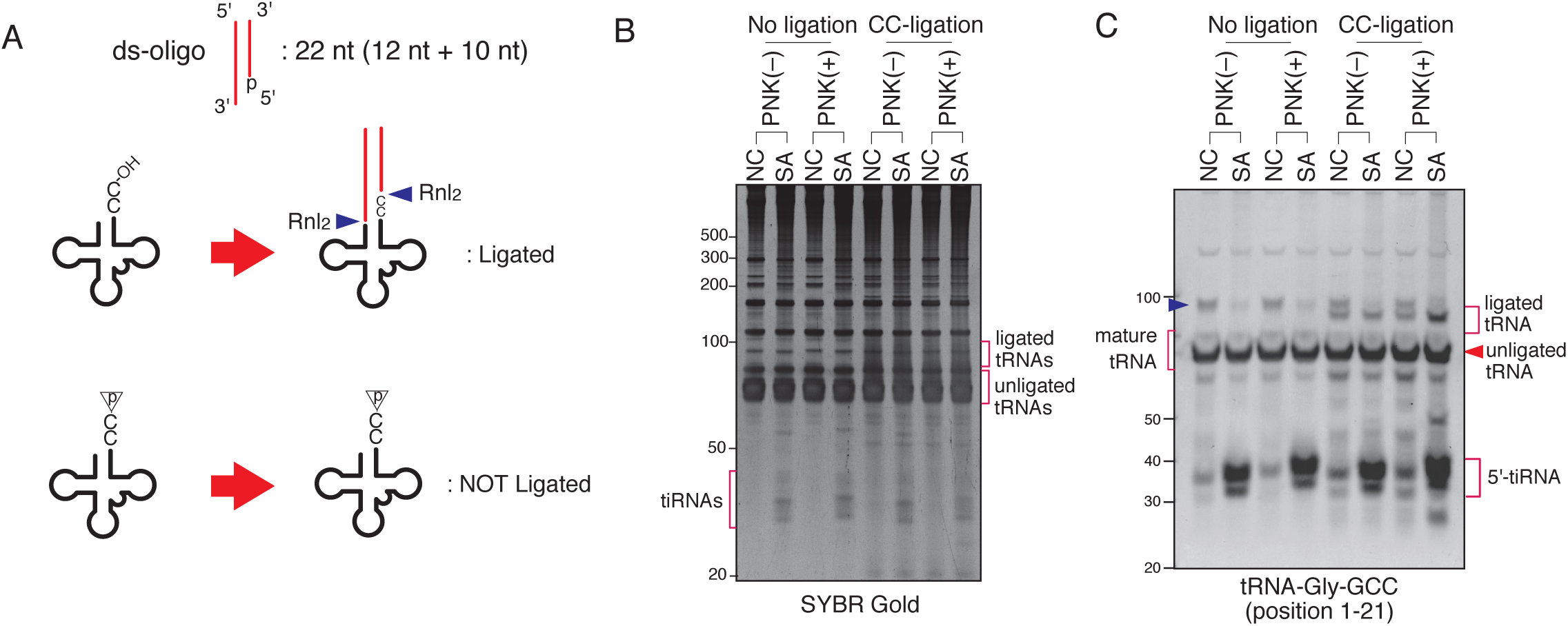
Sodium arsenite treatment induces CC-terminating tRNAs. (A) Schema for CC-specific ligation. Ligation products is generated only when CC-terminating tRNAs have hydroxyl residue on their 3’-end. (B-C) Sodium arsenite treatment increases CC-terminating tRNAs with 2’,3’-cyclic phosphate or 3’-phosphate residue. (B) SYBR Gold staining and (C) Northern blotting of CC-specific ligation products. The blue arrowhead indicates the band for pre-tRNA.

**Figure 6.**
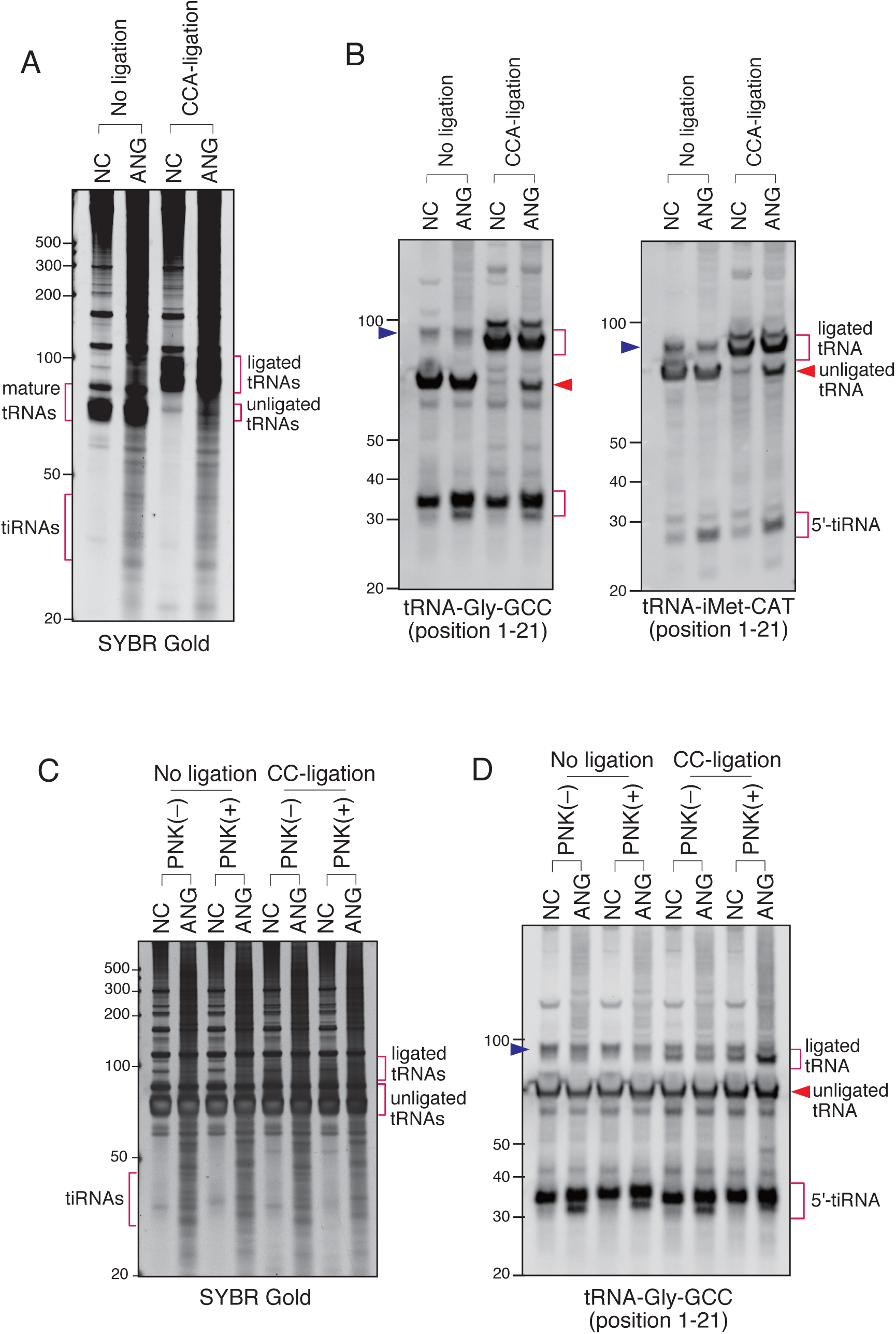
ANG induces CCA-deactivated tRNAs *in vitro*. Total RNAs were subjected to *in vitro* ANG digestion followed by CCA-specific or CC-specific ligation. (A-B) CCA-specific ligation. (A) SYBR Gold staining and (B) Northern blotting. (C-D) CC-specific ligation combined with PNK pretreatment. (C) SYBR Gold staining and (D) Northern blotting. The blue arrowheads indicate the bands for pre-tRNAs.

Our data suggest that SA stress triggers the cleavage of CCA ends of tRNA *in vivo* (Fig. 4 and 6). Although SA treatment readily results in oxidative stress, arsenic is a heavy metal with multiple pleiotropic toxic effects affecting different cellular processes. Treatment of cells with N-acetyl-cysteine (NAC), a potent antioxidant and reactive oxygen species scavenger (44), does not prevent SA-induced CCA cleavage (Fig. S6A-B). Moreover, treatment with high dose of hydrogen peroxide (H_2_O_2_), a potent inducer of oxidative stress (45), also does not trigger CCA cleavage (Fig. 6C-D) in contrast to its previously reported specific effect on tRNA splicing resulting in the accumulation of 5’-leader-exon fragments of tRNA-Tyr-GTA ((46), and Fig. S6D). This suggest that production reactive oxygen species is not a primarily trigger of SA-induced CCA cleavage.

In contrast to ANG, SA triggers cleavage of other RNA substrates (e.g. rRNAs (Fig. 7A)) suggesting that SA activates other RNases. To determine whether ANG is directly involved in the CCA cleavage, we created a U2OS cell line variant with genetic knockout of ANG gene (ΔANG). When WT and ΔANG cell lines were treated with SA, we observed no differences in the CCA cleavage as judged by the CCA-specific ligation of tRNA-Gly-GCC and tRNA-iMet-CAT (Fig. 7B-C) suggesting that ANG does not cleave the CCA-ends of tRNAs *in vivo*. In the same time, we observed that SA also triggers production of tiRNAs in ΔANG U2OS cells suggesting that other cellular RNases are capable to cleave tRNAs in the absence of ANG.

**Figure 7.**
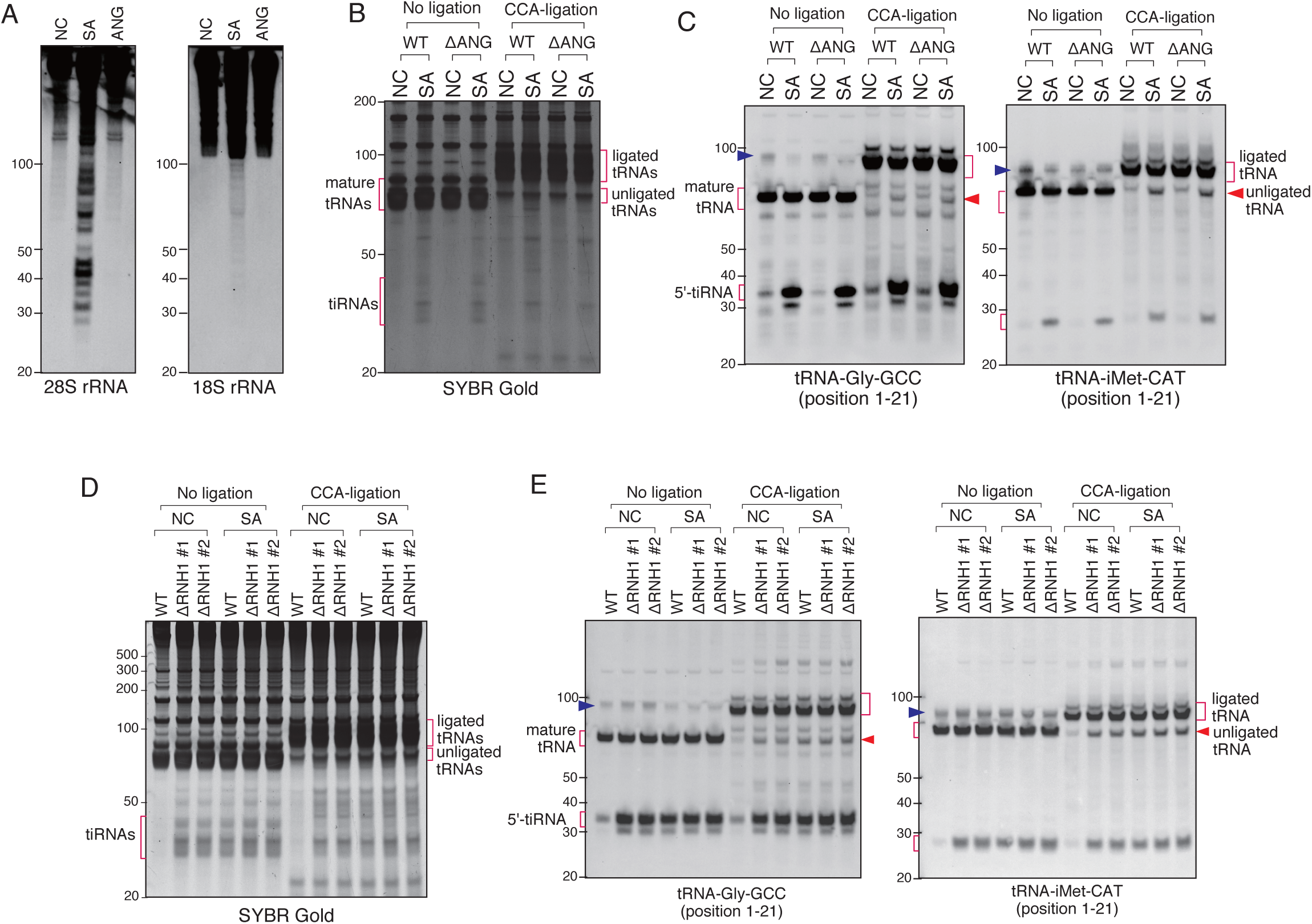
Sodium arsenite-induced CCA-deactivation is through other RNase A family members than ANG. (A) Sodium arsenite treatment induces rRNA fragmentation but ANG does not. (B-C) Sodium arsenite treatment induces CCA-deactivation even in ANG-deleted cells. (B) SYBR Gold staining and (C) Northern blotting of the CCA-ligation products. Sodium arsenite-induced CCA-deactivation is through the activation of RNase A family members. (D-E) Constitutive CCA-deactivation in RNH1-deleted cells. CCA-specific ligation was performed with ΔRNH1 cells with/without sodium arsenite treatment. The blue arrowheads indicate the bands for pre-tRNAs.

RNH1 (Ribonuclease/Angiogenin inhibitor 1) binds and inhibits activities of any RNase belonging to the RNase A superfamily (47). To determine whether a CCA-cleaving enzyme is a member of RNase A superfamily, we created two RNH1 knockout U2OS cell lines (ΔRNH1 #1 and #2). As is seen in Fig. 7D (no ligation, NC), in contrast to WT U2OS cells, ΔRNH1 U2OS cell lines promote spontaneous formation of tiRNAs even in the absence of SA treatment. Under SA treatment, there is no further enhancement of tiRNA production in ΔRNH1 cells (Fig. 7D-E, no ligation) suggesting that in the absence of RNH1, RNase A superfamily members are fully activated and tRNA cleavage is “saturated”. Also, it suggests that SA is unlikely to stimulate tRNA cleavage by other RNases, which do not belong to RNase A superfamily. Importantly, as judged by the CCA-specific ligation assay, in the absence of RNH1, ligation efficiency in ΔRNH1 U2OS cells (in the absence and presence of SA) was similar to the ligation efficiency observed in WT U2OS cells treated with SA (Fig. 7D-E, CCA-ligation).

## DISCUSSION

In this study, we tested whether ANG cleaves the 3’-CCA termini of tRNAs *in vivo*. If ANG efficiently cleaves 3’-CCA termini *in vivo*, it could contribute to stress-induced translational repression by mechanisms distinct from the reported before, where tRNA fragments directly interact with translational machinery (13,48-52). However, our data using RNA-seq reveal that ANG does not affect the percentage of CCA- or CC-terminating species in tRNA fractions, thus demonstrating that ANG does not efficiently cleave 3’-CCA termini *in vivo* (Fig. 1B, Table S3). In addition, we show that more than 98% of 3’-tiRNAs have 3’-CCA after ANG treatment, a percentage that exceed the 90.22% observed under control conditions (Fig. 1C and Table S3). These RNA-seq based data suggest that ANG cleaves CCA termini of tiRNAs to a minimal extent and that it does not precede cleavage at the anticodon loop as suggested by *in vitro* studies (23). In addition, our RNA-seq data show that nearly all tRNAs have intact 3’-CCA termini following ANG cleavage of their anticodon loops *in vivo*.

We acknowledge some limitations to our RNA-seq study. First, in our RNA-seq, we generated the libraries from the tRNA fraction (50-110 nt) or tiRNA fraction (20-50 nt) independently in order to generate enough depth of reads. Therefore, we cannot directly compare the results between the tRNA and tiRNA fractions. For example, we cannot directly analyze the proportion of tRNAs that were cleaved by ANG, or the ratio of anticodon cleavage to 3’-CCA cleavage by ANG. Second, because we defined the reads that overlapped the 3’-ends of tRNA genes as “3’-end containing reads” (Fig S1), some irregular reads could be included such as fragments which had 3’-trailer sequences (e.g. the read with ID:008 in Fig S1C). Such irregular reads could contribute to the decrease in %CCA because they would be categorized as “others”. Because most of the 3’-tiRNAs and tRNAs had CCA at their 3’-termini and were categorized as “CCA-added” (Table S3), such irregular fragments were estimated to be rare.

Therefore, we also validated the differences between ANG-mediated cleavage of tRNAs *in vitro* and *in vivo* using three complimentary biochemical approaches (Fig. S2-3, Fig. 3-5). First approach relies on the RtcB-mediated RNA ligation based on the ability of the enzyme to ligate RNA substrates possessing 2’,3’ cyclic phosphate moiety on their 3’-end (Fig. S2). ANG has the same catalytic properties as RNase A, cleaving the 3’-side of pyrimidine residues to produce a 2’, 3’-cyclic phosphate at the 3’-end of the 5’-cleavage product (e.g., 5’-tiRNA) and a 5’-hydroxyl residue on the 5’-end of the 3’-cleavage product (e.g., 3’-tiRNA). If the CCA triplet is targeted and cleaved by ANG, this would also result in 2’, 3’-cyclic phosphate. In marked contrast to *in vivo* ANG treatment (Fig. S3F-J), when using tiRNA/tRNA fractions generated by *in vitro* ANG digestion (Fig.S3A-E), RtcB ligated not only 5’-tiRNAs but also 3’-tiRNAs and tRNAs to a synthetic oligo (Fig. S3). This demonstrates that ANG can cleave CCA termini of tRNAs *in vitro*, in agreement with previously published data (23).

The second approach depends on the CCA-specific ligation using ligase Rnl2 and ds-oligo in 2’,3’-cyclic phosphate independent manner where Rnl2 is only able to ligate tRNAs with intact CCA ends (Fig. 3). In agreement with RNA-seq data from *in vivo* experiments, there is no difference between ligation efficiency of untreated and ANG-treated tRNAs, although there is a slight, yet reproducible, increase in unligated tRNA signal in the case of SA-treated cells (Figure 4C). Importantly, we developed third ligation approach based on the ability of Rnl2 to ligate only CC-terminating tRNAs with 3’-hydroxyl group. In this method, if an RNase cleaves tRNAs to produce variants terminating with CC-end with cyclic 2’,3’-phosphate, such tRNA substrate will not be ligated (Fig. 5A). Again, and in agreement with other data, ligation of tRNAs from control and SA-treated cells shows that majority of tRNA species are CCA-terminating. Only slight increase in the population of 3’-CC terminating tRNAs with cyclic 2’,3’-phosphate is observed under oxidative stress (Fig. 5B-C). Importantly, SA itself is not capable to promote tRNA/CCA ends cleavage *in vitro* (Fig. S4). In contrast, ANG-mediated digestion of tRNAs *in vitro* promotes CCA cleavage, based on both CCA- and CC-specific ligations (Fig. 6).

It was also suggested that ANG-induced tRNA CCA cleavage may be reversible via repair of the cleaved CCA-ends by the CCA-adding enzyme TRNT1 (23). It should be noted that TRNT1 would need to resolve the cyclic phosphate to a 3’-hydroxyl residue before repair, the activity that has not been attributed to enzyme. Theoretically, the 2’, 3’-cyclic nucleotide 3’-phosphodiesterase (CNPase) could be involved with this reaction. However, although it has been suggested that CNPase is involved with the metabolism of endogenous 2’, 3’-cAMP especially in the myelin sheath (53), the involvement of the enzyme in tRNA metabolism including tRNA splicing has not been established in the vertebrates (54). Our data suggest that efficient depletion of TRNT1 (Fig. 2A) does not change percentage of CCA- and CC-terminating tiRNAs derived from cytosolic tRNAs in cells treated with recombinant ANG (Fig. 2D and Table S4). At the same time, TRNT1 knockdown significantly decreases population of CCA-terminating tiRNAs originating from mitochondrial tRNA-Ser-GCT, in agreement with its susceptibility to TRNT1 hypofunction (37). Thus, although capable of targeting both mitochondrial and cytosolic tRNAs, ANG cleaves them in the anticodon loops.

Finally, experiments using genetic knockouts of ANG and RNH1 suggest that: 1) SA-induced CCA cleavage is independent of ANG (Fig. 7B-C); 2) RNH1 regulates SA-induced CCA cleavage (Fig. 7D-E) suggesting that an unidentified RNase is a member of the RNase A superfamily; 3) RNH1 negatively regulates production of tiRNAs (Fig. 7D-E); 4) In the absence of ANG, other ribonuclease(s) can generate tiRNAs (Fig. 7B-C). The latter observation is in agreement with the recent publication from the Dutta lab that also show that tiRNAs can also be generated by unknown RNase(s) in the conditions of high concentration SA-induced stress (55). Although their data suggest that overexpression of ANG leads to cleavage of specific tRNA subset, our data suggest that treatment with recombinant ANG leads to production of 5’-tiRNAs from all tRNAs (Fig. 1D). Nonetheless, both studies and our previous work (36) agree that ANG is a stress-responsive RNase capable of tiRNA production but also other RNases can play a role in stress-induced tRNA cleavage, and that some tRNA fragments (like tRNA-Gly-GCC) are present in cells also in unstressed conditions. Whether different RNases act in stress-specific or cell-specific manner to generate tiRNAs as suggested in tRNA microarray studies (36) needs to be determined. Interestingly, SA promotes tRNA cleavage in oxidative stress-independent manner *in vivo* (Fig. S6) suggesting that it may activate downstream signalling pathways that in turn may regulate activity of specific RNases. Future studies on tRNA cleavage will determine details of biogenesis of various tRNA fragments and roles that various RNases play in this process.

## SUPPLEMENTARY DATA

Supplementary Data are available

## ACKNOWLEDGEMENT

We thank Victoria Ivanova and Dhwani Dave for assistance with preliminary experiments and all Anderson and Ivanov lab members for helpful critiques.

## REFERENCES

1. Strydom, D.J., Fett, J.W., Lobb, R.R., Alderman, E.M., Bethune, J.L., Riordan, J.F. and Vallee, B.L. (1985) Amino acid sequence of human tumor derived angiogenin. Biochemistry, 24, 5486–5494.

2. Lee, F.S. and Vallee, B.L. (1989) Characterization of ribonucleolytic activity of angiogenin towards tRNA. Biochemical and biophysical research communications, 161, 121–126.

3. Shapiro, R. and Vallee, B.L. (1989) Site-directed mutagenesis of histidine-13 and histidine-114 of human angiogenin. Alanine derivatives inhibit angiogenin-induced angiogenesis. Biochemistry, 28, 7401–7408.

4. Shapiro, R., Riordan, J.F. and Vallee, B.L. (1986) Characteristic ribonucleolytic activity of human angiogenin. Biochemistry, 25, 3527–3532.

5. Curran, T.P., Shapiro, R. and Riordan, J.F. (1993) Alteration of the enzymatic specificity of human angiogenin by site-directed mutagenesis. Biochemistry, 32, 2307–2313.

6. Sheng, J. and Xu, Z. (2016) Three decades of research on angiogenin: a review and perspective. Acta Biochim Biophys Sin (Shanghai), 48, 399–410.

7. Pereira, E.R., Liao, N., Neale, G.A. and Hendershot, L.M. (2010) Transcriptional and post-transcriptional regulation of proangiogenic factors by the unfolded protein response. PloS one, 5.

8. Kishimoto, K., Yoshida, S., Ibaragi, S., Yoshioka, N., Okui, T., Hu, G.F. and Sasaki, A. (2012) Hypoxia-induced up-regulation of angiogenin, besides VEGF, is related to progression of oral cancer. Oral Oncol, 48, 1120–1127.

9. Lyons, S.M., Fay, M.M., Akiyama, Y., Anderson, P.J. and Ivanov, P. (2017) RNA biology of angiogenin: Current state and perspectives. RNA Biol, 14, 171–178.

10. Pizzo, E., Sarcinelli, C., Sheng, J., Fusco, S., Formiggini, F., Netti, P., Yu, W., D’Alessio, G. and Hu, G.F. (2013) Ribonuclease/angiogenin inhibitor 1 regulates stress-induced subcellular localization of angiogenin to control growth and survival. J Cell Sci, 126, 4308–4319.

11. Yamasaki, S., Ivanov, P., Hu, G.F. and Anderson, P. (2009) Angiogenin cleaves tRNA and promotes stress-induced translational repression. The Journal of cell biology, 185, 35–42.

12. Fu, H., Feng, J., Liu, Q., Sun, F., Tie, Y., Zhu, J., Xing, R., Sun, Z. and Zheng, X. (2009) Stress induces tRNA cleavage by angiogenin in mammalian cells. FEBS letters, 583, 437–442.

13. Ivanov, P., Emara, M.M., Villen, J., Gygi, S.P. and Anderson, P. (2011) Angiogenin-induced tRNA fragments inhibit translation initiation. Mol Cell, 43, 613–623.

14. Panas, M.D., Ivanov, P. and Anderson, P. (2016) Mechanistic insights into mammalian stress granule dynamics. The Journal of cell biology, 215, 313–323.

15. Kedersha, N., Ivanov, P. and Anderson, P. (2013) Stress granules and cell signaling: more than just a passing phase? Trends in biochemical sciences, 38, 494–506.

16. Emara, M.M., Ivanov, P., Hickman, T., Dawra, N., Tisdale, S., Kedersha, N., Hu, G.F. and Anderson, P. (2010) Angiogenin-induced tRNA-derived stress-induced RNAs promote stress-induced stress granule assembly. The Journal of biological chemistry, 285, 10959–10968.

17. Anderson, P., Kedersha, N. and Ivanov, P. (2015) Stress granules, P-bodies and cancer. Biochimica et biophysica acta, 1849, 861–870.

18. Saikia, M., Jobava, R., Parisien, M., Putnam, A., Krokowski, D., Gao, X.H., Guan, B.J., Yuan, Y., Jankowsky, E., Feng, Z. et al. (2014) Angiogenin-cleaved tRNA halves interact with cytochrome c, protecting cells from apoptosis during osmotic stress. Mol Cell Biol, 34, 2450–2463.

19. Rybak, S.M. and Vallee, B.L. (1988) Base cleavage specificity of angiogenin with Saccharomyces cerevisiae and Escherichia coli 5S RNAs. Biochemistry, 27, 2288–2294.

20. Li, Z., Ender, C., Meister, G., Moore, P.S., Chang, Y. and John, B. (2012) Extensive terminal and asymmetric processing of small RNAs from rRNAs, snoRNAs, snRNAs, and tRNAs. Nucleic Acids Res, 40, 6787–6799.

21. Saxena, S.K., Rybak, S.M., Davey, R.T., Jr., Youle, R.J. and Ackerman, E.J. (1992) Angiogenin is a cytotoxic, tRNA-specific ribonuclease in the RNase A superfamily. The Journal of biological chemistry, 267, 21982–21986.

22. Hoang, T.T. and Raines, R.T. (2017) Molecular basis for the autonomous promotion of cell proliferation by angiogenin. Nucleic Acids Res, 45, 818–831.

23. Czech, A., Wende, S., Morl, M., Pan, T. and Ignatova, Z. (2013) Reversible and rapid transfer-RNA deactivation as a mechanism of translational repression in stress. PLoS Genet, 9, e1003767.

24. Chakraborty, P.K., Schmitz-Abe, K., Kennedy, E.K., Mamady, H., Naas, T., Durie, D., Campagna, D.R., Lau, A., Sendamarai, A.K., Wiseman, D.H. et al. (2014) Mutations in TRNT1 cause congenital sideroblastic anemia with immunodeficiency, fevers, and developmental delay (SIFD). Blood, 124, 2867–2871.

25. Wu, D., Yu, W., Kishikawa, H., Folkerth, R.D., Iafrate, A.J., Shen, Y., Xin, W., Sims, K. and Hu, G.F. (2007) Angiogenin loss-of-function mutations in amyotrophic lateral sclerosis. Ann Neurol, 62, 609–617.

26. Pall, G.S. and Hamilton, A.J. (2008) Improved northern blot method for enhanced detection of small RNA. Nat Protoc, 3, 1077–1084.

27. Lyons, S.M., Achorn, C., Kedersha, N.L., Anderson, P.J. and Ivanov, P. (2016) YB-1 regulates tiRNA-induced Stress Granule formation but not translational repression. Nucleic Acids Res, 44, 6949–6960.

28. Martin, M. (2011) Cutadapt removes adapter sequences from high-throughput sequencing reads. EMBnet journal, 17, 10–12.

29. Karolchik, D., Baertsch, R., Diekhans, M., Furey, T.S., Hinrichs, A., Lu, Y.T., Roskin, K.M., Schwartz, M., Sugnet, C.W., Thomas, D.J. et al. (2003) The UCSC Genome Browser Database. Nucleic Acids Res, 31, 51–54.

30. Langmead, B., Trapnell, C., Pop, M. and Salzberg, S.L. (2009) Ultrafast and memory-efficient alignment of short DNA sequences to the human genome. Genome Biol, 10, R25.

31. Li, H., Handsaker, B., Wysoker, A., Fennell, T., Ruan, J., Homer, N., Marth, G., Abecasis, G., Durbin, R. and Genome Project Data Processing, S. (2009) The Sequence Alignment/Map format and SAMtools. Bioinformatics, 25, 2078–2079.

32. Chan, P.P. and Lowe, T.M. (2009) GtRNAdb: a database of transfer RNA genes detected in genomic sequence. Nucleic Acids Res, 37, D93–97.

33. Putz, J., Dupuis, B., Sissler, M. and Florentz, C. (2007) Mamit-tRNA, a database of mammalian mitochondrial tRNA primary and secondary structures. RNA, 13, 1184–1190.

34. Hu, G., Xu, C. and Riordan, J.F. (2000) Human angiogenin is rapidly translocated to the nucleus of human umbilical vein endothelial cells and binds to DNA. Journal of cellular biochemistry, 76, 452–462.

35. Yang, W. (2011) Nucleases: diversity of structure, function and mechanism. Q Rev Biophys, 44, 1–93.

36. Saikia, M., Krokowski, D., Guan, B.J., Ivanov, P., Parisien, M., Hu, G.F., Anderson, P., Pan, T. and Hatzoglou, M. (2012) Genome-wide identification and quantitative analysis of cleaved tRNA fragments induced by cellular stress. The Journal of biological chemistry, 287, 42708–42725.

37. Sasarman, F., Thiffault, I., Weraarpachai, W., Salomon, S., Maftei, C., Gauthier, J., Ellazam, B., Webb, N., Antonicka, H., Janer, A. et al. (2015) The 3’ addition of CCA to mitochondrial tRNASer(AGY) is specifically impaired in patients with mutations in the tRNA nucleotidyl transferase TRNT1. Hum Mol Genet, 24, 2841–2847.

38. Honda, S., Morichika, K. and Kirino, Y. (2016) Selective amplification and sequencing of cyclic phosphate-containing RNAs by the cP-RNA-seq method. Nat Protoc, 11, 476–489.

39. Honda, S., Loher, P., Shigematsu, M., Palazzo, J.P., Suzuki, R., Imoto, I., Rigoutsos, I. and Kirino, Y. (2015) Sex hormone-dependent tRNA halves enhance cell proliferation in breast and prostate cancers. Proceedings of the National Academy of Sciences of the United States of America, 112, E3816–3825.

40. Tanaka, N., Chakravarty, A.K., Maughan, B. and Shuman, S. (2011) Novel mechanism of RNA repair by RtcB via sequential 2’,3’-cyclic phosphodiesterase and 3’-Phosphate/5’-hydroxyl ligation reactions. The Journal of biological chemistry, 286, 43134–43143.

41. Bullard, D.R. and Bowater, R.P. (2006) Direct comparison of nick-joining activity of the nucleic acid ligases from bacteriophage T4. The Biochemical journal, 398, 135–144.

42. Dittmar, K.A., Goodenbour, J.M. and Pan, T. (2006) Tissue-specific differences in human transfer RNA expression. PLoS Genet, 2, e221.

43. Shigematsu, M., Honda, S., Loher, P., Telonis, A.G., Rigoutsos, I. and Kirino, Y. (2017) YAMAT-seq: an efficient method for high-throughput sequencing of mature transfer RNAs. Nucleic Acids Res, 45, e70.

44. c tt Aldini, G., Altomare, A., Baron, G., Vistoli, G., Carini, M., Borsani, L. and Sergio, F. (2018) N-Acetylcysteine as an antioxidant and disulphide breaking agent: the reasons why. Free Radic Res, 52, 751–762.

45. Sies, H., Berndt, C. and Jones, D.P. (2017) Oxidative Stress. Annu Rev Biochem, 86, 715–748.

46. Hanada, T., Weitzer, S., Mair, B., Bernreuther, C., Wainger, B.J., Ichida, J., Hanada, R., Orthofer, M., Cronin, S.J., Komnenovic, V. et al. (2013) CLP1 links tRNA metabolism to progressive motor-neuron loss. Nature, 495, 474–480.

47. Dickson, K.A., Haigis, M.C. and Raines, R.T. (2005) Ribonuclease inhibitor: structure and function. Prog Nucleic Acid Res Mol Biol, 80, 349–374.

48. Sobala, A. and Hutvagner, G. (2013) Small RNAs derived from the 5’ end of tRNA can inhibit protein translation in human cells. RNA Biol, 10, 553–563.

49. Mleczko, A.M., Celichowski, P. and Bakowska-Zywicka, K. (2018) Transfer RNA-derived fragments target and regulate ribosome-associated aminoacyl-transfer RNA synthetases. Biochim Biophys Acta Gene Regul Mech.

50. Lyons, S.M., Gudanis, D., Coyne, S.M., Gdaniec, Z. and Ivanov, P. (2017) Identification of functional tetramolecular RNA G-quadruplexes derived from transfer RNAs. Nat Commun, 8, 1127.

51. Ivanov, P., O’Day, E., Emara, M.M., Wagner, G., Lieberman, J. and Anderson, P. (2014) G-quadruplex structures contribute to the neuroprotective effects of angiogenin-induced tRNA fragments. Proceedings of the National Academy of Sciences of the United States of America, 111, 18201–18206.

52. Fricker, R., Brogli, R., Luidalepp, H., Wyss, L., Fasnacht, M., Joss, O., Zywicki, M., Helm, M., Schneider, A., Cristodero, M. et al. (2019) A tRNA half modulates translation as stress response in Trypanosoma brucei. Nat Commun, 10, 118.

53. Verrier, J.D., Jackson, T.C., Bansal, R., Kochanek, P.M., Puccio, A.M., Okonkwo, D.O. and Jackson, E.K. (2012) The brain in vivo expresses the 2’,3’-cAMP-adenosine pathway. Journal of neurochemistry, 122, 115–125.

54. Myllykoski, M., Seidel, L., Muruganandam, G., Raasakka, A., Torda, A.E. and Kursula, P. (2016) Structural and functional evolution of 2’,3’-cyclic nucleotide 3’-phosphodiesterase. Brain research, 1641, 64–78.

55. Su, Z., Kuscu, C., Malik, A., Shibata, E. and Dutta, A. (2019) Angiogenin generates specific stress-induced tRNA halves and is not involved in tRF-3-mediated gene silencing. The Journal of biological chemistry.

