## Supplementary Information for "Multiple ribonuclease A family members cleave transfer RNAs in response to stress"

#### Supplementary Method

##### Purification of recombinant *E.coli* RtcB

The open reading frame of *E. coli* RtcB was cloned into pET28a vector and transformed into BL21 (DE3) *E. coli*. Cultures derived from single kanamycin-resistant transformants were grown at 37 °C until the $A^{600}$ reached 0.6–0.8. Then protein expression was induced with 0.1 mM IPTG (isopropyl-β-D-thiogalactopyranoside). Incubation was continued at room temperature for 16 hours with shaking. His6-RtcB protein was purified with TALON metal affinity resin (Clontech) according to the manufacturer's protocol. His6-tag was removed with Thrombin (GE Healthcare) during overnight dialysis, and thrombin was removed with Benzamidine Sepharose beads (GE Healthcare).

##### *In vitro* RtcB RNA ligation assay

5'-hydroxyl, 3'-biotinylated RNA oligo was synthesized by Integrated DNA Technology. The sequence of the oligo is:

5'-uuggguguagcucagugguagagcgcgugcu/biotin/-3'.

10 µg total RNA from ANG-treated U2OS cells or 25 µg total RNA treated with ANG *in vitro* were electrophoresed on 15% Urea-TBE gel, and 5'-tiRNA, 3'-tiRNA and tRNA fractions were gel-purified as RNA substrates in 16 µl of nuclease-free water. 4 µL and 6 µL were subjected to SYBR Gold staining and ligation reaction, respectively. For ligation reaction, 0.5 µM RNA oligo and RNA substrates were incubated in 20 µL reaction mixtures containing 50 mM Tris-HCl (pH 7.4), 2 mM MnCl, 100 µM GTP, 2 U/µL RNasin (Promega) and 1 µM RtcB, for 30 min at 37 °C. After purification using Direct-zol RNA Microprep (Zymo Research), RNAs were electrophoresed on 15% Urea-TBE gel and then transferred to Hybond N+ nylon membrane (Amersham Pharmacia Biotech). The oligo-ligated products were

detected by SYBR Gold (Thermo Fisher Scientific) and HRP-conjugated streptavidin (Jackson ImmunoResearch) coupled with ECL.

**Supplementary Figures and Tables Legends**

Supplementary Figure 1. Workflow for preparation of RNA sequencing library. (A) Flowchart of the preparation of the library. OH- hydroxyl group, P- 5'-phosphate, P in the triangle – cyclic 2'-3'-phosphate. (B) Diagram of trimming for the calculation of CCA - or CC-terminating reads. (C) Extraction of 3'-end containing reads and calculation of the proportion of CCA- or CC-terminating reads. In this example, %CCA and %CC are 57.1% (4/7) and 14.3% (1/7), respectively. (D) Calculation of the proportion of CCA-added (%CCA) or CC-terminating (%CC) fragments. The proportions were calculated as follows: %CCA = 100 x (A+D+G) / (X+Y+Z), %CC = 100 x (B+E+H) / (X+Y+Z). Note that all 22 genes ending with CC have C before CC (i.e. all the genes end with "CCC"). Therefore, the mature tRNAs derived from these genes should end with CCCCCA.

Supplementary Figure 2. Diagram of ligation products of the reaction with Rtcb. The RNA oligo is ligated to only 5'-products generated by ANG-mediated cleavage because of the existence of 2', 3'-cyclic phosphate residue on their 3'-ends. (a) 5'-tiRNAs are ligated to the oligo and 5'-tiRNA-oligo products will be generated. (b) Intact (i.e. CCA-added) 3'-tiRNAs are not ligated to the oligo. (c) If CCA-termini of 3'-tiRNAs are cleaved by ANG, 3'-tiRNA-oligo products will be generated by RtcB. (d) Intact (i.e. CCA-added) tRNAs are not ligated to the oligo. (e) If CCA-termini of tRNAs are cleaved by ANG, tRNA-oligo products will be generated.

Supplementary Figure 3. Evaluation of ANG cleavage specificity towards tRNA anticodon loops and 3'-CCA ends using RtcB-based *in vitro* ligation assay. (A)-(E) Ligation reactions with tiRNA/tRNA fractions derived from *in vitro* ANG digestion. (A) Representative SYBR Gold staining of gel-purified tiRNAs. Synthetic 5'-hydroxyl, 3'-

biotinylated RNA oligo (oligo, 31 nt) used for ligation reaction is shown. (B) SYBR Gold staining of the ligation products derived from tiRNA fractions isolated from *in vitro* ANG cleavage. Ligation products are indicated by red triangles. Note that the bands around 30 nt are unreacted oligo (31 nt). (C) Detection of ligation products (B) by streptavidin-biotin system. Note that both 5'-tiRNA and 3'-tiRNA were ligated to the oligo and ligation products were detected. ND: not detected. (D) SYBR Gold staining of the reacted tRNA fraction. Gel-purified tRNA fraction without ligation reaction is shown as control. Ligation products between the control oligo and ANG-digested tRNA fraction are indicated by red triangle. (E) Detection of ligation products by streptavidin-biotin system. A band around 100 nt (indicated by red triangle) was detected in the RtcB reaction between the oligo and ANG-digested tRNA fraction, in contrast to the reaction with tRNAs without ANG digestion. (F)-(J) Ligation assays with tiRNA/tRNA fraction derived from ANG-treated U2OS cells. (F) Representative SYBR Gold staining of gel-purified tiRNAs from U2OS cells. Synthetic 5'-hydroxyl, 3'-biotinylated RNA oligo (oligo, 31 nt) used for ligation reaction is shown. (G) SYBR Gold staining of the ligation products derived from the ligation of 5'- and 3'- tiRNA fractions with control oligo, or from the ligation between 5'- and 3'-tiRNAs. (H) Detection of ligation products (G) by streptavidin-biotin system. (I) SYBR Gold staining of the reacted tRNA fraction. Gel-purified tRNA fraction without ligation reaction is also shown. (J) RtcB did not generate any ligation product from the oligo and tRNA fraction. (K) Summary of the experiments (A-J). Estimated length was calculated for tRNA-Gly-GCC (72 nt). Seventy-two nucleotides is the most common length of tRNA genes (186 genes out of 631 genes). Mature tRNA-Gly-GCC (75 nt) is cleaved by ANG into 5'-tiRNA (34 nt) and 3'-tiRNA (41 nt). See also Supplementary Figure 3 for details.

Supplementary Figure 4. Sodium arsenite does not induce any RNA degradation *in vitro*. Total RNAs from U2OS cells were incubated with sodium arsenite at the concentrations as indicated at room temperature for 1 hour. (A) SYBR Gold staining and (B) Northern blotting.

Supplementary Figure 5. Genotype of ΔANG and ΔRNH1 cells. (A-C) Genotype of ΔANG cells. (A) Sequencing of ANG genomic locus. Initiator ATG is highlighted in green. (B) Predicted protein product. N-terminal signal peptide is underlined, and amino acids downstream of Cas9-induced insertion are highlighted in red. (C) Western blotting confirming the loss of ANG expression. ACTB was used as a loading control. (D-H) Genotype of ΔRNH1 cells. (D) Sequencing of ΔRNH1 genomic locus in clone#1. Initiator ATG and terminal codon are highlighted in green and red, respectively. (E) Predicted protein product in ΔRNH1 clone#1 cells. (F) Sequencing of ΔRNH1 genomic locus in clone#2. (G) Predicted protein product in ΔRNH1 clone#2 cells. (G) Western blotting confirming the loss of RNH1 expression. ACTB was used as a loading control.

Supplementary Figure 6. Sodium arsenite-induced CCA-deactivation is independent of oxidative stress. (A-B) N-acetylcysteine(NAC) pre-treatment has no effect on sodium arsenite-induced CCA-deactivation. (A) SYBR Gold staining and (B) Northern blotting. U2OS cells were treated with 20 mM NAC (pH 7.5) (Sigma-Aldrich) for 1 hr before sodium arsenite treatment, followed by RNA purification and CCA-specific ligation. NAC pre-treatment did not decrease the amount of CCA-deactivated tRNAs, in contrast to slight decrease of tiRNAs. (C-D) Hydrogen peroxide ($H_2O_2$) does not induce CCA-deactivation. (C) SYBR Gold staining and (D) Northern blotting. U2OS cells were treated with 500 μM of $H_2O_2$ for 1 hr, followed by RNA purification and CCA-specific ligation. $H_2O_2$ treatment did not induce either tiRNA production or CCA-deactivation. On the other hand, $H_2O_2$ treatment induced the generation of the 5'-leader-exon fragment derived from tRNA-Tyr-GTA as previously reported (1).

Supplementary Table 1. The sequences of oligos for CCA-specific or CC-specific ligation method. N: mix of A, T, G or C. n: mix of a, u, g or c. 5Phos: 5'-phosphorylated. 3Bio: 3'-biotinylated.

Supplementary Table 2. Sequences of probes for Northern blotting.

Supplementary Table 3. Effect of ANG treatment on CCA-termini of (A) tRNAs and (B) 3-tiRNAs. Values are means ± SE (n=3).

Supplementary Table 4. TRNT1 does not affect the proportion of CCA-added 3'-tiRNAs in ANG-treated cells. (A) Effect of TRNT1 knockdown on the proportion of CCA-added 3'-tiRNAs derived from mitochondrial tRNA-Ser-GCT. Values are means ± SE (n=3). *: $p<0.05$ VS siControl-NC, ** $p<0.01$ VS siControl-NC, †: $p<0.01$ VS siTRNT1-NC, ††: $p<0.01$ VS siTRNT1-NC. (B) Effect of TRNT1 knockdown on the proportion of CCA-added and CC-added 3'-tiRNAs in ANG-treated cells. Values are means ± SE (n=3). **: $p<0.01$ VS siControl-NC, ††: $p<0.01$ VS siControl-ANG

**Reference**

1. Hanada, T., Weitzer, S., Mair, B., Bernreuther, C., Wainger, B.J., Ichida, J., Hanada, R., Orthofer, M., Cronin, S.J., Komnenovic, V. *et al.* (2013) CLP1 links tRNA metabolism to progressive motor-neuron loss. *Nature*, **495**, 474-480.

### Supplementary Figure 1.

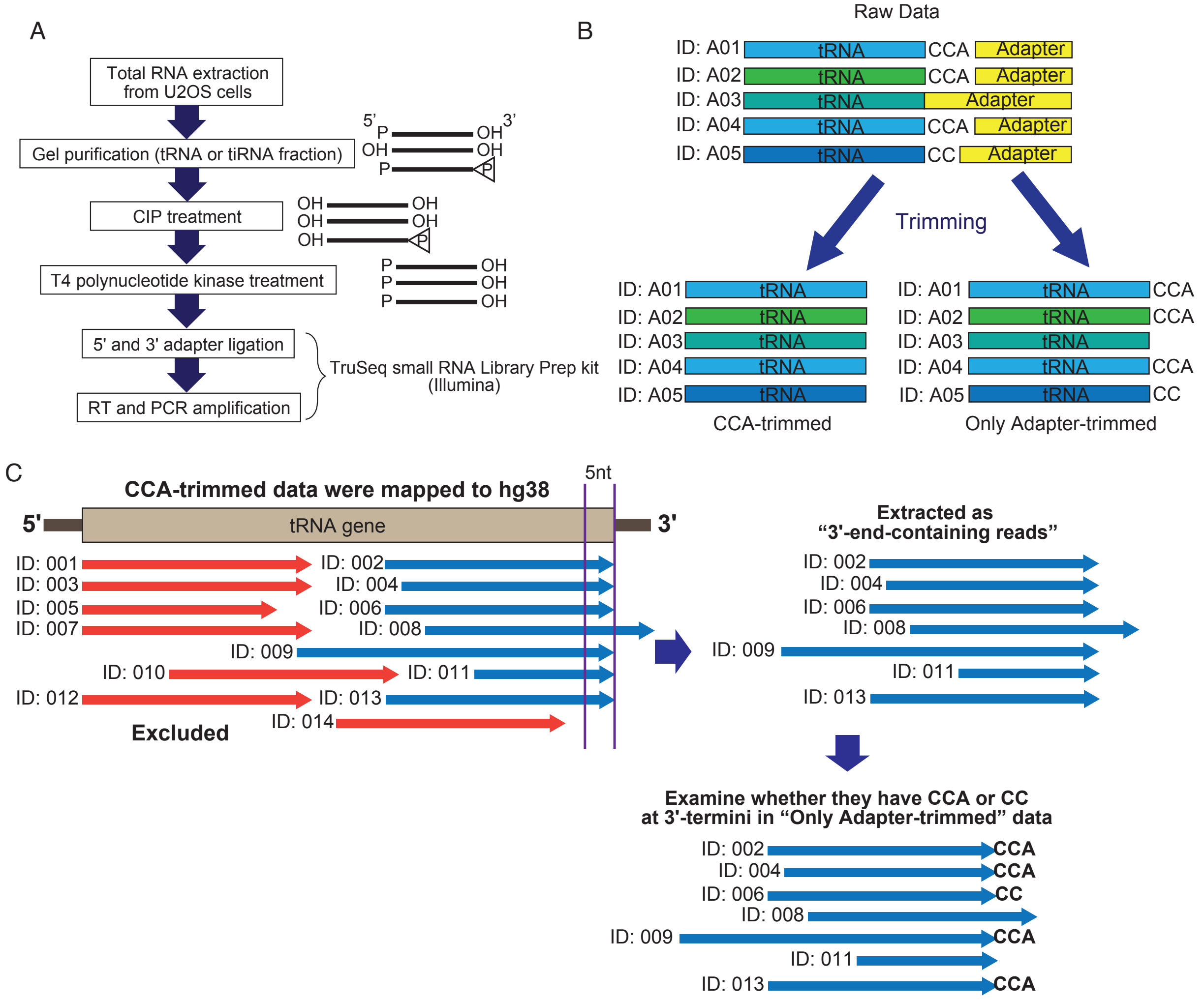


D

| | tRNA genes ending with CCA | tRNA genes ending with (C)CC | the other tRNA genes | total |
|---|---|---|---|---|
| | 109 genes | 22 genes | 500 genes | 631 genes |
| CCA-added | A: ending with "CCACCA" | D: ending with "CCCCCA" | G: ending with "CCA" | A+D+G |
| CC-terminating | B: ending with "CCACC" | E: ending with "CCCCC" | H: ending with "CC" | B+E+H |
| the others | X-(A+B) | Y-(D+E) | Z-(G+H) | X+Y+Z-(A+B+D+E +G+H) |
| total counts | X | Y | Z | X+Y+Z |

### Supplementary Figure 2.

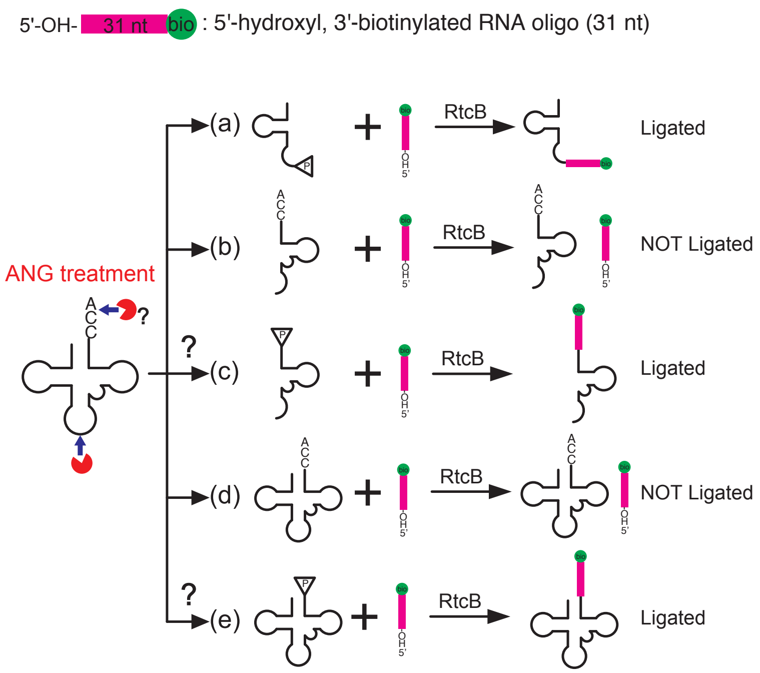

### Supplementary Figure 3.

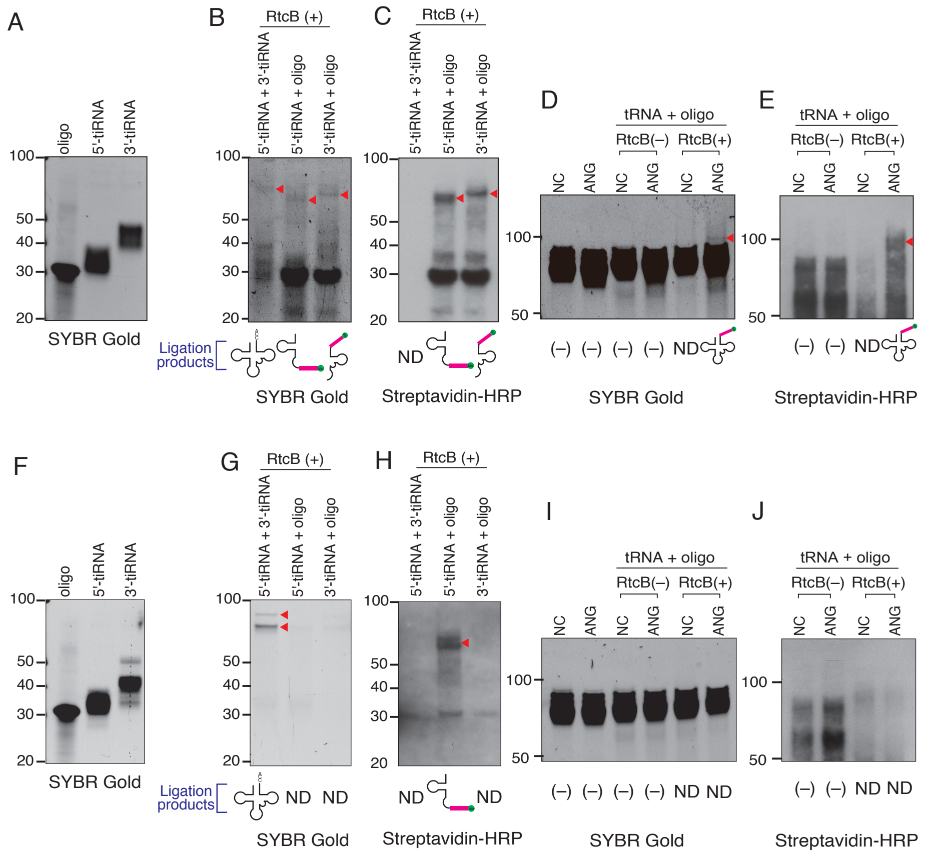


K

| Ligation reaction | Estimated length | Ligation Product | |
|---|---|---|---|
| | | *in vitro* ANG digestion | *in vivo* ANG treatment |
| 5'-tiRNA + 3'-tiRNA | 75 nt | Repaired tRNA | Repaired tRNA |
| 5'-tiRNA + oligo | 65 nt | Ligated | Ligated |
| 3'-tiRNA + oligo | 72 nt | Ligated | No ligation product |
| tRNA + oligo | 105 nt | Ligated | No ligation product |

### Supplementary Figure 4.

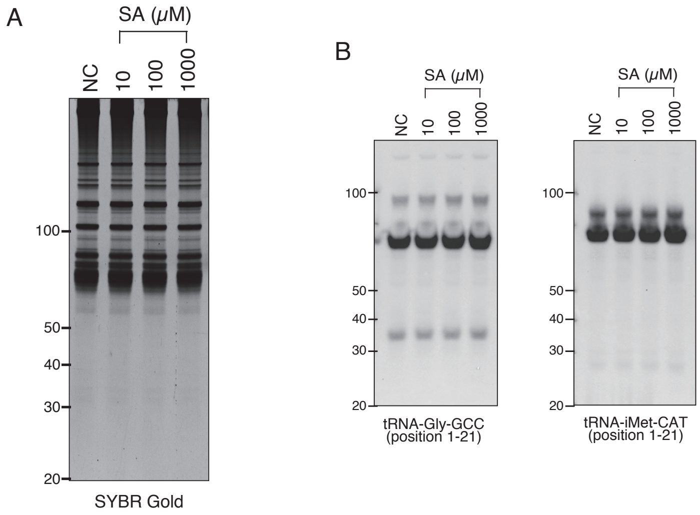

### Supplementary Figure 5.

A

```
WT     GGAAGAGATGGTGATGGGCCTGGGCGTTTTGTTGTTGGTCTTCG-TGCTGGGTCTGGGTCTGACCCCACCGACCCTGGCTCAGGA
ΔANG   GGAAGAGATGGTGATGGGCCTGGGCGTTTTGTTGTTGGTCTTCGTTGCTGGGTCTGGGTCTGACCCCACCGACCCTGGCTCAGGA
       ******************************************** ****************************************
```

B

**Wild Type ANG**

MVMGLGVLLLVFVLGLGLTPPTLAQDNSRYTHFLTQHYDAKPQGRDDRYCESIM
RRRGLTSPCKDINTFIHGNKRSIKAICENKNGNPHRENLRISKSSFQVTTCKLHG
GSPWPPCQYRATAGFRNVVVACENGLPVHLDQSIFRRP*

**ΔANG**

MVMGLGVLLLVFVAGSGSDPTDPGSG*

C

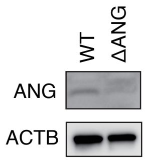


D

```
ΔRNH1 #1 Allele 1   CACCATGAGCCTGGACATCCAGAGCCTGGACATCCAGTTGTGAGGAGCTGAGCGACGCTAGATGGGCCGA
ΔRNH1 #1 Allele 2   CACCATGAGCCTGGACATCCAGAGC---------------------------------CTAGATGGGCCGA
RefSeq              CACCATGAGCCTGGACATCCAGAGCCTGGACATCCAG-TGTGAGGAGCTGAGCGACGCTAGATGGGCCGA
                    *************************                                 *************
```

E

**Wild Type RNH1**

MSLDIQSLDIQCEELSDARWAELLPLLQQCQVVRLDDCGLTEARCKDISSALRVNPALAELNLRSNELGDVGVHCVLQG
LQTPSCKIQKLSLQNCCLTGAGCGVLSSTLRTLPTLQELHLSDNLLGDAGLQLLCEGLLDPQCRLEKLQLEYCSLSAAS
CEPLASVLRAKPDFKELTVSNNDINEAGVRVLCQGLKDSPCQLEALKLESCGVTSDNCRDLCGIVASKASLRELALGSN
KLGDVGMAELCPGLLHPSSRLRTLWIWECGITAKGCGDLCRVLRAKESLKELSLAGNELGDEGARLLCETLLEPGCQLE
SLWVKSCSFTAACCSHFSSVLAQNRFLLELQISNNRLEDAGVRELCQGLGQPGSVLRVLWLADCDVSDSSCSSLAATLL
ANHSLRELDLSNNCLGDAGILQLVESVRQPGCLLEQLVLYDIYWSEEMEDRLQALEKDKPSLRVIS

**ΔRNH1 #1 Allele 1**

MSLDIQSLDIQL*

**ΔRNH1 #1 Allele 2**

MSLDIQS*

F

```
ΔRNH1 #2   CACCATGAGCCTGGACATCCAGAGC---------------------------------CTAGATGGGCCGA
RefSeq     CACCATGAGCCTGGACATCCAGAGCCTGGACATCCAG-TGTGAGGAGCTGAGCGACGCTAGATGGGCCGA
           *************************                                 *************
```

G

**ΔRNH1 #2**

MSLDIQS*

H

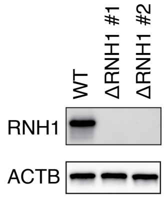

### Supplementary Figure 6.

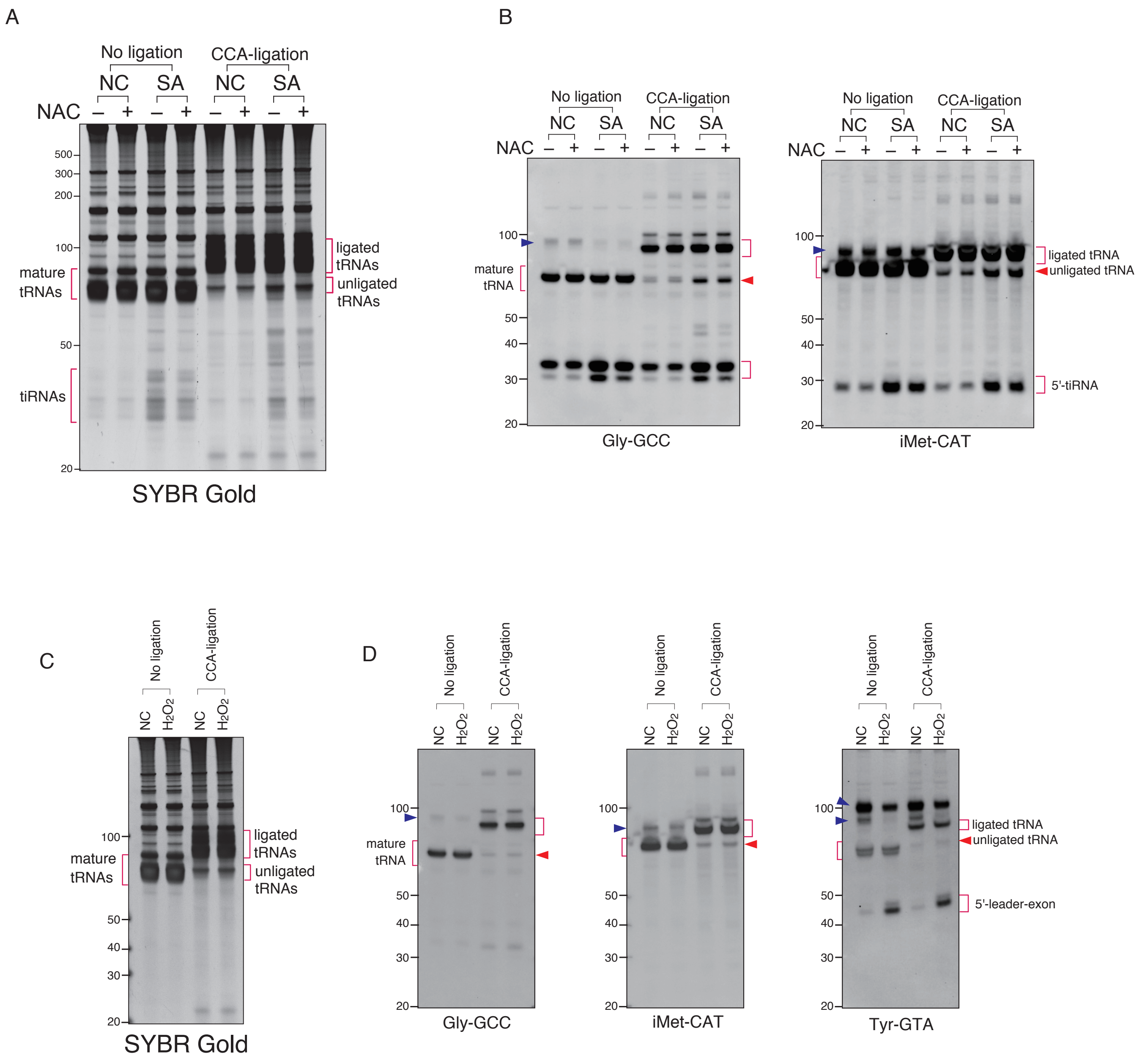

### Supplementary Table 1.

| Oligos | | Sequence |
|---|---|---|
| Hairpin-oligo for CCA-specific ligation | | /5Phos/cgcacugcTTTTTGCAGTGCGTGGN |
| double-strand oligo for CCA-specific ligation | 5'-oligo | ACTGGATACTGgn |
| | 3'-oligo | /5Phos/GTATCCAGTT/3Bio/ |
| double-strand oligo for CC-specific ligation | 5'-oligo | ACTGGATACggn |
| | 3'-oligo | /5Phos/GTATCCAGTT |

### Supplementary Table 2.

| Probes | Position | Sequences |
|---|---|---|
| tRNA-Ala-AGC | 1-32 | 5'-AAGCACGCGCTCTACCACTGAGCTACACCCCC-3' |
| tRNA-Arg-ACG | 1-21 | 5'-TATCCATTGCGCCACTGGCCC-3' |
| tRNA-Asn-GTT | 1-33 | 5'-GCCGAACGCGCTAACCGATTGCGCCACAGAGAC-3' |
| tRNA-Asp-GTC | 1-34 | 5'-CAGGCGGGGATACTCACCACTATACTAACGAGGA-3' |
| tRNA-Cys-GCA | 1-32 | 5'-AGTCAAATGCTCTACCACTGAGCTATACCCCC-3' |
| tRNA-Gln-CTG | 1-29 | 5'-CAGAGTGCTAACCATTACACCATGGAACC-3' |
| tRNA-Glu-CTC | 1-21 | 5'-TAACCACTAGACCACCAGGGA-3' |
| tRNA-Gly-GCC | 1-21 | 5'-CTACCACTGAACCACCCATGC-3' |
| tRNA-His-GTG | 1-31 | 5'-CGCAGAGTACTAACCACTATACGATCACGGC-3' |
| tRNA-Ile-AAT | 1-33 | 5'-GCACCACGCTCTAACCAACTGAGCTAACCGGCC-3' |
| tRNA-Leu-CAG | 1-21 | 5'-AGACCGCTCGGCCATCCTGAC-3' |
| tRNA-Lys-CTT | 1-21 | 5'-TACCGACTGAGCTAGCCGGGC-3' |
| tRNA-Met-CAT | 1-31 | 5'-ACTGACGCGCTACCTACTGCGCTAACGAGGC-3' |
| tRNA-iMet-CAT | 1-21 | 5'-CTTCCGCTGCGCCACTCTGCT-3' |
| tRNA-Phe-GAA | 1-32 | 5'-GTCTAACGCTCTCCCAACTGAGCTATTTCGGC-3' |
| tRNA-Pro-TGG | 1-21 | 5'-ATACCCCTAGACCAACGAGCC-3' |
| tRNA-SeC-TCA | 1-34 | 5'-GCCTGCACCCCAGACCACTGAGGATCATCCGGGC-3' |
| tRNA-Ser-GCT | 1-21 | 5'-TAACCACTCGGCCACCTCGTC-3' |
| tRNA-Thr-AGT | 1-33 | 5'-GACAGGCGCTTTAACCAACTAAGCCACGGCGCC-3' |
| tRNA-Trp-CCA | 1-31 | 5'-GTCAGACGCGCTGCCGTTGCGCCACGAGGTC-3' |
| tRNA-Tyr-GTA | 1-34 | 5'-CAGTCCTCCGCTCTACCAACTGAGCTATCGAAGG-3' |
| tRNA-Val-AAC | 1-32 | 5'-GGCGAACGTGATAACCACTACACTACGGAAAC-3' |
| 28S rRNA | | 5'-GGGTGAACAATCCAACGCTTGGTG-3' |
| 18S rRNA | | 5'-AGAGGAGCGAGCGACCAAAGGAA-3' |

### Supplementary Table 3.

A

| tRNA-fraction | Control | ANG | P value |
|---|---|---|---|
| total fragments | 188,990 ± 7,551 | 164,810 ± 3,691 | 0.0452 |
| CCA-added | 179,947 ± 7,085 | 156,044 ± 3,600 | 0.0396 |
| CC-terminating | 1,170 ± 163 | 1,020 ± 89 | 0.4663 |
| the others | 7,873 ± 524 | 7,745 ± 451 | 0.8623 |
| %CCA-added | 95.22 ± 0.22 | 94.68 ± 0.22 | 0.1578 |
| %CC-terminating | 0.61 ± 0.06 | 0.62 ± 0.05 | 0.9433 |

B

| tiRNA-fraction | Control | ANG | P value |
|---|---|---|---|
| total fragments | 38,375 ± 2,787 | 240,173 ± 27,604 | 0.0019 |
| CCA-added | 34,621 ± 2,539 | 236,797 ± 27,353 | 0.0018 |
| CC-terminating | 1,151 ± 301 | 1,765 ± 270 | 0.2039 |
| the others | 2,602 ± 410 | 1,161 ± 12 | 0.0729 |
| %CCA-added | 90.22 ± 0.81 | 98.58 ± 0.08 | 0.0005 |
| %CC-terminating | 2.93 ± 0.55 | 0.73 ± 0.05 | 0.0166 |

### Supplementary Table 4.

A

| tiRNA fraction of mito-tRNA-Ser-GCT | NC | | ANG | |
|---|---|---|---|---|
| | siControl | siTRNT1 | siControl | siTRNT1 |
| total fragments | 59.0 ± 10.0 | 38.4 ± 3.5 | 21.4 ± 4.0 | 7.2 ± 0.6 |
| CCA-added | 36.6 ± 7.4 | 17.5 ± 1.9* | 13.2 ± 2.5 | 2.8 ± 0.5 |
| CC-terminating | 11.0 ± 0.8 | 9.3 ± 0.2 | 4.0 ± 0.8 | 2.3 ± 0.3 |
| the others | 11.4 ± 2.2 | 11.6 ± 1.6 | 4.2 ± 1.1 | 2.0 ± 0.1 |
| %CCA-added | 61.4 ± 2.0 | 45.5 ± 1.7** | 61.5 ± 6.7 | 39.0 ± 1.0†† |
| %CC-terminating | 19.4 ± 2.5 | 24.5 ± 1.7 | 19.1 ± 2.6 | 32.1 ± 1.5† |

B

| tiRNA fraction | NC | | ANG | |
|---|---|---|---|---|
| | siControl | siTRNT1 | siControl | siTRNT1 |
| total fragments | 22,306 ± 628 | 25,342 ± 1,176 | 124,504 ± 16,783** | 109,038 ± 5,698‡‡ |
| CCA-added | 19,728 ± 529 | 22,233 ± 1,119 | 122,073 ± 16,579** | 106,604 ± 5,531‡‡ |
| CC-terminating | 664 ± 22 | 796 ± 43 | 934 ± 83 | 860 ± 89 |
| the others | 1,913 ± 96 | 2,313 ± 109 | 1,496 ± 95 | 1,574 ± 91‡‡ |
| %CCA-added | 88.45 ± 0.26 | 87.70 ± 0.58 | 98.98 ± 0.27** | 97.77 ± 0.06‡‡ |
| %CC-terminating | 2.98 ± 0.01 | 3.15 ± 0.23 | 0.77 ± 0.10** | 0.79 ± 0.05‡‡ |
